# Real-time Image Denoising of Mixed Poisson-Gaussian Noise in Fluorescence Microscopy Images using ImageJ

**DOI:** 10.1101/2021.11.10.468102

**Authors:** Varun Mannam, Yide Zhang, Yinhao Zhu, Evan Nichols, Qingfei Wang, Vignesh Sundaresan, Siyuan Zhang, Cody Smith, Paul W Bohn, Scott Howard

## Abstract

Fluorescence microscopy imaging speed is fundamentally limited by the measurement signal-to-noise ratio (SNR). To improve image SNR for a given image acquisition rate, computational denoising techniques can be used to suppress noise. However, common techniques to estimate a denoised image from a single frame are either computationally expensive or rely on simple noise statistical models. These models assume Poisson or Gaussian noise statistics, which are not appropriate for many fluorescence microscopy applications that contain quantum shot noise and electronic Johnson–Nyquist noise, therefore a mixture of Poisson and Gaussian noise. In this paper, we show convolutional neural networks (CNNs) trained on mixed Poisson and Gaussian noise images to overcome the limitations of existing image denoising methods. The trained CNN is presented as an open-source ImageJ plugin that performs real-time image denoising (within tens of milliseconds) with superior performance (SNR improvement) compared to the conventional fluorescence microscopy denoising methods. The method is validated on external datasets with out-of-distribution noise, contrast, structure, and imaging modalities from the training data and consistently achieves high performance (> 8 dB) denoising in less time than other fluorescence microscopy denoising methods.

## 1 Introduction

In modern biology, fluorescence microscopy plays a vital role in functional and structural imaging [1]. However, the imaging speed of fluorescence microscopy is limited by the inherent noise in the system [2]. Fluctuations in the photon emission events contribute to quantum noiseshot noise (from the stream of photons) which follows Poisson statistics per time period. Additionally, microscope electronic components contribute Johnson–Nyquist noise (or thermal noise), which follows a Gaussian distribution per time period. The combination of both of these contributions leads to fluorescence microscopy systems containing a mixed Poisson-Gaussian (MPG) noise. Due to this inherent noise in the system, the acquired image has a limited signal-to-noise ratio (SNR) that hinders the underlined ground truth information about the biological sample. Therefore, a robust denoising approach is required to address these two distinct noise distributions. In fluorescence microscopy, the statistics of each noise source affect images differently. To illustrate these differences, we simulate the effects of Gaussian noise and Poisson noise individually on a clean image in Figure 1. The clean image represents bovine pulmonary artery endothelial cells (BPAE, Invitrogen FluoCells slide#1 F36924, mitochondria labeled with MitoTracker Red CMXRos, F-actin labeled with Alexa Fluor 488 phalloidin, nuclei labeled with DAPI) imaged with a commercial fluorescence microscope (Nikon A1R-MP laser scanning confocal microscope equipped with a Nikon Apo LWD 40x, 1.15 NA water immersion objective and more details are provided in the Supplementary Note S1) with a 4 mW excitation power and 50 frame averaging to produce a high SNR image. Gaussian noise and Poisson noise are added to the clean image (SNR of 25 dB with reference to the clean image) and shown as “With Gaussian noise,” and “With Poisson noise,” images, respectively. The “Clean image” is obtained by averaging 50 noisy images in the same field of view (FOV). Here, the Gaussian noise is independent of clean image pixel intensity, whereas the Poisson noise is pixel intensity-dependent. We can observe this behaviour in the marked ROI center region, i.e., the smooth gray area (Nuclei region). Hence, Poisson noise and Gaussian noise affect the clean image differently, which makes the MPG noise denoising a challenging task.

**Figure 1:**
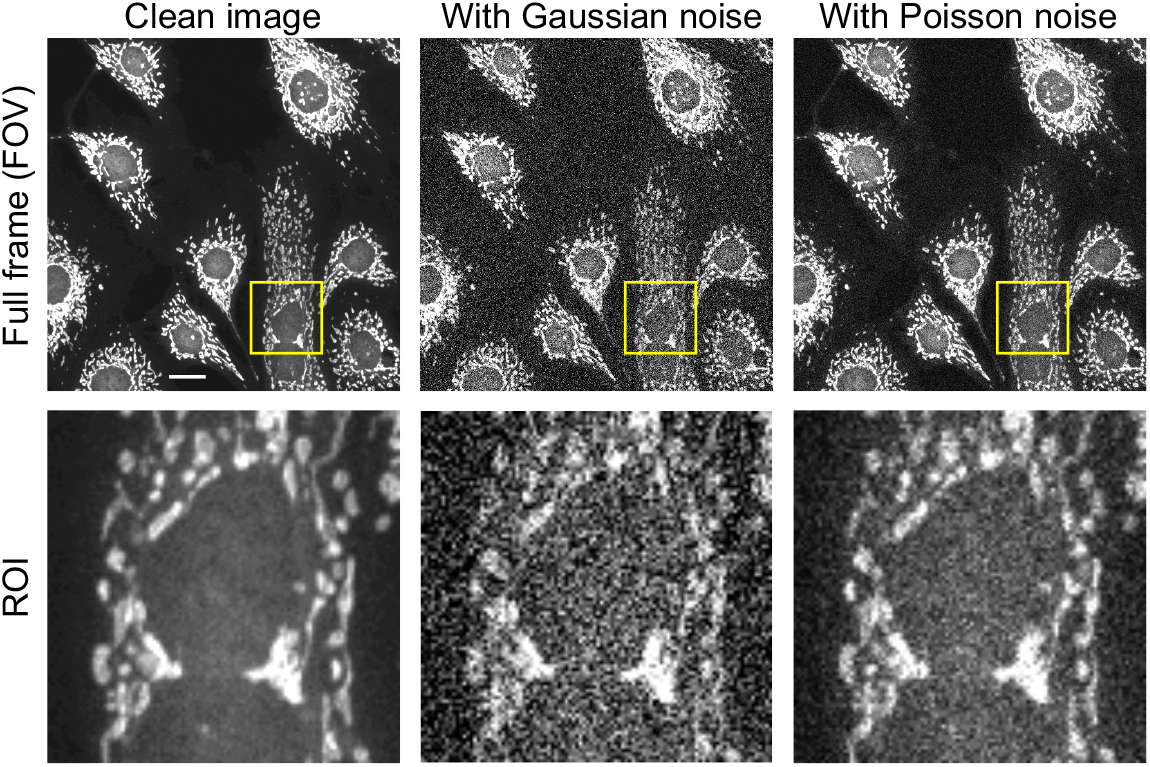
Illustration of different types of noise with a BPAE cell sample. The top row indicates the full frame images and the PSNR of the synthetic noisy image is 25 dB with reference to clean image (obtained by averaging 50 noisy images in the same FOV). The bottom row shows the selected region of interest (ROI) as shown in the yellow box of the corresponding top row images. Full frame and ROI dimensions are 512 ×512 and 125 ×125 respectively. Scale bar: 20 *µ*m.

To improve fluorescence microscopy signal-to-noise ratio (SNR), one can increase the sample dosage by either increasing the excitation power or the pixel dwell time. However, this method could lead to photobleaching or photodamage and cannot be used for imaging real-time dynamics in living animals. Furthermore, lower dosage leads to noisy (with low SNR) images and thereby posing a challenge for applications like cell classification and segmentation [3]. Denoising algorithms are available to improve image SNR, but are either designed specifically for Gaussian or Poisson noise (thereby not suitable for the other distribution or MPG noise), or are capable of denoising MPG noise but require long computation time to achieve relatively small SNR improvement. Therefore, there is a need for a computational fast single-frame denoising method that works for all experimental noise distributions: Poisson, Gaussian, and MPG.

Reiterating the mathematical notation of different noise distributions using 1^*st*^ and 2^*nd*^ order moments (or simply mean and variance) [4], the Gaussian noise distribution is represented as standard normal distribution *n*_*g*_ *∼ 𝒩* (0, *b*) where the mean is zero and the variance is *b*. Similarly, the Poisson noise distribution can be represented as *n*_*p*_ *∼ 𝒩* (*y*_*i*_, *ay*_*i*_) where the true pixel value (*y*_*i*_) is the mean and Poisson variance is *ay*_*i*_ (conversion factor or quantum efficiency of *a* times *y*_*i*_). Assuming that the Poisson and Gaussian processes are independent, the MPG noise distribution is represented as *n*_mixed_ ∼ 𝒩 (*y*_*i*_, *ay*_*i*_ + *b*) where the mean is the true pixel value (*y*_*i*_) and the variance is the addition of two independent noise variances (*ay*_*i*_ + *b*). Hence, MPG noise distribution is similar to Poisson noise with an additional Gaussian noise variance (*b*).

In addition to the above-described noise statistics, fluorescence microscopy systems using detector arrays (e.g., charge-coupled device (CCD) or complementary metal-oxide semiconductor (CMOS)) can experience a “fixed-pattern noise” due to differences in detector responsivity and dark-current between pixels. Such noise commonly appears as horizontal or vertical “stripes” in low-signal regions of an image. The signal variance in the presence of fixed-pattern noise also follows the form *ay*_*i*_ + *b*, but with *a* and *b* varying at each pixel. Denoising such systems using calibration or noise statistics are possible [5], but requires prior details on each specific system. Fixed pattern noise can also be addressed using photon event centroid algorithm [6], flat-field correction method [7] and automatic correction of CMOS-related noise using camera properties combined with sparse filtering [5], or calibration by measuring the pixel-specific dark current and gain, then compensating for it via subtraction and division, respectively. In this paper, we will primarily consider images that do not experience fixed-pattern noise or have compensated for fixed-patter noise via calibration.

In order to increase the SNR of an image, additional information must be included to estimate the original, denoised image. If the noise distribution is known *a priori* to be Gaussian, simple filtering (by assuming noise exceeds signal at high spatial frequencies by using mean, median, bilateral filters) or machine-learning based approaches such as non-local means (NLM) [8], block matching 3D filtering (BM3D) [9], K-means singular value decomposition (K-SVD) [10], expected patch log-likelihood (EPLL) [11], and weighted nuclear norm minimization (WNNM) [12] can all be used to leverage knowledge of the noise distribution and to leverage information from similar features within the same image. If the noise distribution is known to be Poisson, the above methods can be used by (1) converting the Poisson noise into Gaussian noise using a variance stabilization transform (VST) (e.g., generalized Anscombe transform (GAT [13])), (2) denoising using popular Gaussian denoising algorithms, and (3) converting the denoised results back using an inverse VST [13]. Such popular Poisson denoising methods include Poisson NLM (PNLM) [14], non-local principle component analysis (NL-PCA) [15], VST+NLM and VST+BM3D which apply VST, NLM or BM3D Gaussian denoising methods, and inverse VST, sequentially, on an image. If the noise is from a MPG distribution, more complicated methods are needed to estimate the relative contributions of each process at each pixel (e.g., PURE-LET [16]).

Overall, each of these image denoising methods have trade-offs between denoising performance, computation speed, and applicability to a specific noise distribution. In this paper, we show how a convolutional neural network (CNN) trained on mixed Poisson-Gaussian noise can in fact achieve high-performance denoising, quickly, on images with Poisson and/or Gaussian noise distributions.

Recently, CNN based fluorescence microscopy image denoising as demonstrated improved denoising performance and speed [17]. CNN-based image denoising methods can be classified into two categories: supervised and self-supervised. Supervised methods require training the network on a dataset prior to performing denoising on experimental images and include content-aware image restoration (CARE) [17], denoising convolutional neural networks (DnCNN) [18], and Noise2Noise (N2N) [19]. Self-supervised methods train the network on the same image it is performing denoising on and Noise2Void (N2V) [20], probabilistic Noise2Void (PN2V) [21], Noise2Fast [22], and other methods [23, 24]. Supervised CNN training methods typically provide higher performance than the self-supervised methods due to the inclusion of additional information in target images. In some conditions, self-supervised denoising methods can yield very comparable performance to supervised methods [19]. In this paper, we utilize a supervised CNN model that is capable of removing MPG noise from a single frame nearly in real-time (within a few ms). By producing and utilizing a large, multiple species, and multiple microscope modality training dataset [25], the trained model outperforms existing image denoising methods. More importantly, our method demonstrates superior denoising performance compared to the existing image denoising approaches including ML-based methods independent of the noise distribution (either Poisson or Gaussian or MPG noise) presented in most of the fluorescence microscopy images.

To address the fundamental problem of denoising microscopy images containing MPG noise sources, we train and evaluate well-known CNN models on MPG noise microscopy data from the FMD dataset [26] in Sections 2 and 3. In Section 4, the trained ML model is validated by evaluating performance in a diverse set of microscopy images, including the W2S fluorescence microscopy dataset [27], with out-of-training-distribution SNR and contrast. The method consistently produces high-quality denoised images and is packaged as an ImageJ plugin to reduce the barrier-to-entry for applying denoising CNNs to microscopy data.

## 2 Methods

As described in Section 1, the fundamental stochastic nature of MPG noise requires introducing some *a priori* knowledge to the denoising process. Analytical approaches use knowledge of the statistical properties of noise, but are computationally slow and not well suited for MPG sources. Deep CNNs, however, are data-driven and expressive enough to overcome the complex statistical nature of MPG noise. To perform denoising using CNNs, a network architecture and training datasets must be chosen. In this paper, we choose to focus on the Noise2Noise [19] and DnCNN network [18] architectures for two reasons. First, they have demonstrated superior performance to other CNN architectures [20]. Second, robust performance in diverse microscopy environments requires training on large datasets. Supervised training of the Noise2Noise and DnCNN models can be achieved in reasonable time (≈ 4 hours for the entire FMD dataset on a single Nvidia 1080-ti GPU) while training other networks (e.g., Noise2Void) on the dataset can take a prohibitively long time (≈ 3.50 hours for a single image training on the GPU). Noise2Void is a popular denoising method that is capable of self-supervised learning, so we will therefore compare supervised trained model performance to self-supervised Noise2Void performance in Section 4.4.1. We chose to train these two models on our FMD dataset [25, 26] that contains 12,000 raw noisy images from confocal [28], two-photon [29], and widefield [30] microscopes. The FMD dataset also helps to benchmark various denoising techniques on the fluorescence images. The creation of the training and testing datasets from the FMD dataset is explained in the Supplementary Note S1.

The first ML model we evaluate is based on a deep CNN with the Noise2Noise architecture [19] trained to denoise fluorescence images with MPG noise. The ML model is similar to the originally published Noise2Noise architecture [19] with the addition of a non-linear activation (tanh) layer after the final convolutional layer [26], adaptive learning (one cycle policy) during the training phase, and carefully tuned hyper-parameters such as batch size and learning rate for achieving best fluorescence microscopy image denoising performance. Here onwards we named the modified model as “Noise2Noise plugin”.

In the Noise2Noise plugin CNN model, the input and target are both noisy images within the same FOV (which is the case when the ground truth is absent or its extraction is difficult). This ML model is beneficial compared to our second ML model which is based on the DnCNN architecture [18] that requires ground truth since getting the clean image (typically acquired with the increased dosage) is usually difficult, especially when imaging living species. Figure 2 shows the complete Noise2Noise plugin architecture as our first ML model. Here we divide the ML architecture into five blocks. Block A to C represent an encoder structure, and Block C to E represent a decoder structure. In the encoder, the image feature size (256 × 256) is reduced by selecting only essential features of the image (with more channels to select different features), and at block C, the images with feature map size (8 × 8) are less noisy when compared to the noisy input image. In the decoder, this noise-free image is restored to the original noisy image feature size (256 × 256). In this work, both input and target are noisy images. At the *i*-th pixel of the input and target images, the noisy measurements are *z*_1_ = *y* + *n*_1_ and *z*_2_ = *y* + *n*_2_, where *y* is the real value and *n*_1_ and *n*_2_ are MPG noise values corresponding to the input and target, respectively. The Noise2Noise plugin model estimates the real value *y* instead of the noisy target *z*_2_ since *n*_2_ is random. The Noise2Noise plugin model can predict the clean image since training happens on the noisy image pair as input and target and this ML model works well as long as the clean image is the average of noisy target images within a FOV. The other ML model is based on a deep CNN with the DnCNN architecture trained to denoise the fluorescence images with MPG noise. The DnCNN ML model can estimate the residual (only noise) for a given noisy input image and the denoised image is obtained by subtracting the estimated residual from the noisy input image. Unlike the supervised DnCNN ML model which requires clean images as the target, the Noise2Noise plugin ML model utilizes the noisy image as the target. The training and testing of the proposed ML models are explained in the Supplementary Note S2.

**Figure 2:**
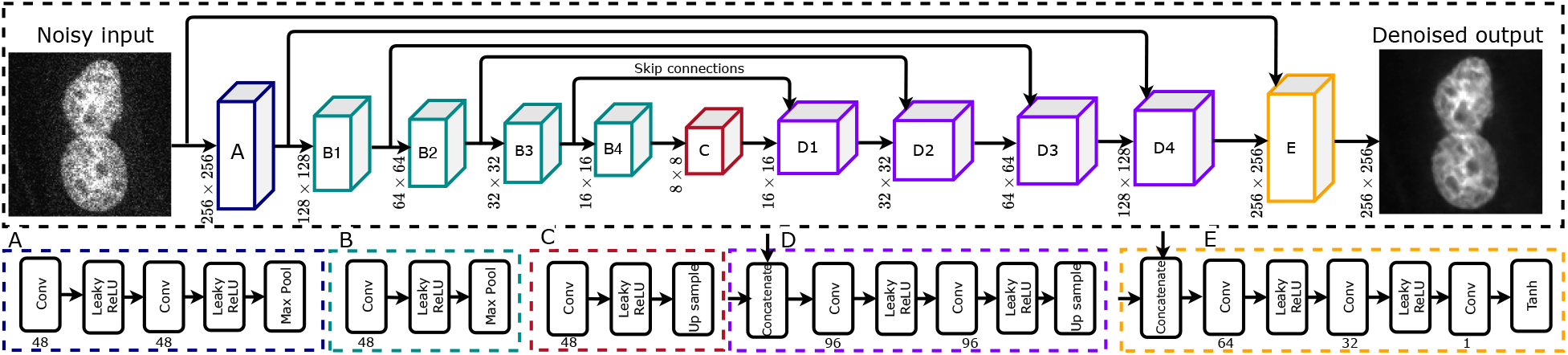
The Noise2Noise plugin architecture we use to train on the FMD dataset. We divide the ML model mainly into two sections, namely encoder (from block A to C: contrast path) and decoder (from block C to E: expansion path). At every block, input dimensions are provided, and the corresponding layers in each block are represented in the sub-blocks along with the number of output channels in the convolution provided in **bold**. Skip connections are useful for adding features from encoder to decoder which might be suppressed in the latent space (at C block). Noisy input and Denoised output image dimensions are 256 × 256.

Interestingly, both the Noise2Noise and DnCNN in literature use relatively simple U-Net and fully convolutional neural networks, respectively, yet achieve high-quality denoising results. However, more advanced architectures and methods are commonly required to perform microscopy image identification and segmentation [31, 32, 33, 34]. While the model capacity of simple U-Net and convolutional neural networks is reduced compared to advanced methods, denoising models only require training a model to discern between the presence and absence of general common features while image identification and segmentation models must be trained to discriminate between specific high-detail features. We therefore employ simple, yet high-performance, denoising networks in this study and evaluate performance on a variety of image samples in Section 4.

While the proposed ML approach is fast and accurate, the complexity of implementing the ML model is a barrier to entry. To address this barrier, we integrated our ML models as plugins to ImageJ, one of the most widely used biomedical image processing packages [35]. Two ImageJ plugins are presented for two ML models: Noise2Noise plugin and DnCNN plugin. We have developed our ImageJ plugins based on the ImageJ-tensorflow library [36]. Here the library provides the minimal set of functions in ImageJ to load the pre-trained ML model weights as a graph and perform a set of convolutional layer operations on the input image (loaded by ImageJ) which results in the output of the trained ML model. To accommodate more image formats, we have extended the ImageJ-tensorflow library to handle 16-bit and 32-bit image data-types (in addition to the original 8-bit images) for both training and inference. The required image pre-processing (e.g., pixel value normalization) is performed automatically by the plugin, allowing users to load and process images without manually pre-processing. Both ML models were trained in Keras with TensorFlow as the backend. ImageJ plugins simplify applying denoising CNNs by integrating into biomedical researcher’s existing workflow. Additionally, the plugins support graphical processing unit (GPU) computing, resulting in a significant speed increase compared to conventional ImageJ image denoising plugins [37, 17, 38] that run on a central processing unit (CPU). Also, recent implementation methods such as DeepImageJ plugin [38] and ZeroCost4Microscopy Jupyter python code [39] can be used with our pre-trained ML model weights (saved in .zip format) to perform image denoising. The ImageJ plugin design and code are explained in the Supplementary Note S5. The following section describes performance and analysis of the trained denoising ML models when applied to mixed Poisson-Gaussian noise in fluorescence microscopy images.

## 3 Results and Discussion

The Noise2Noise plugin ML model is trained with different initial learning rates (ILRs). The training and testing loss (mean square error) are provided in the Supplementary Note S2. We evaluate the trained ML model performance on test-mix data by selecting four images in each microscopy modality (48 images), as seen in Figure 3.

**Figure 3:**
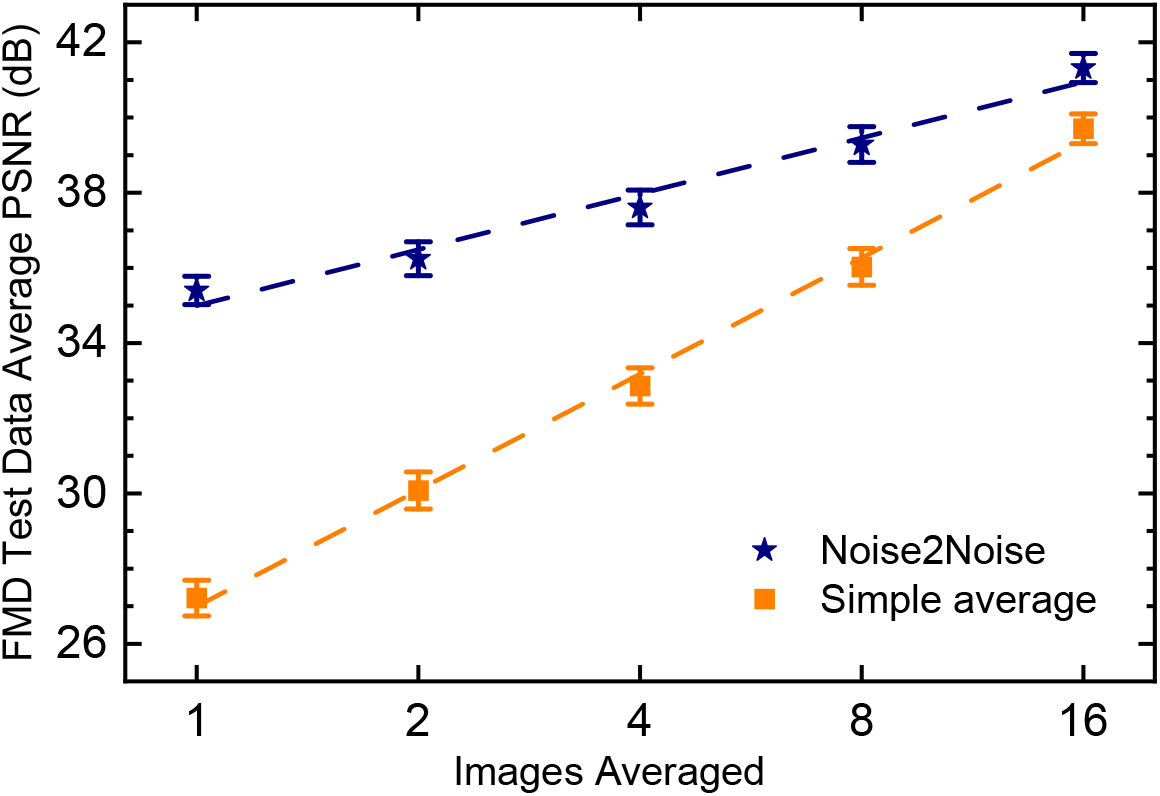
Average PSNR of the test-mix dataset which contains 48 images of the MPG noise samples from different microscopes in the FMD dataset. Here the error bars indicate one standard error below and above the average value, respectively.

To quantify image SNR, we evaluate peak SNR (PSNR) of an image relative to a clean version of the image. PSNR is the measure of mean square error (MSE) between two images (MSE) normalized to the peak value in an image so that MSE between images with different bit depths or signal levels can be compared. PSNR of a given (*X*) with reference to ground truth image (*Y*) in the same FOV is defined as 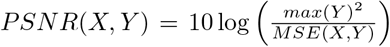, where 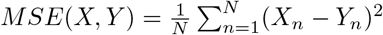 is the average mean-square error of *X* and *Y* with *N* pixels. Results show an average PSNR improvement of **8.13 dB** in the denoised images compared to the average PSNR of the raw images (Images Averaged = 1) in the test-mix data.

For deeper understanding along with more detailed analysis on these results, we perform a simulation that maps the fluorescence microscopy modality and corresponding noise characteristics of the acquired image by estimating the noise variance parameters (*a* and *b*) using the algorithm provided in [40] on the FMD dataset. The results are tabulated in Table 1.

**Table 1:** Estimation of noise parameters (*a* and *b*) using the FMD dataset. The estimated *a* and *b* are average values of raw noisy images from 20 different FOVs (50 images within each FOV). CLSM: confocal laser scanning microscopy; TPEF: two-photon excitation microscopy; WF: widefield microscopy.

| Modality | Sample | Estimated $a$ | Estimated $b$ |
| --- | --- | --- | --- |
| CLSM | BPAE (Nuclei) | $1.76 \times 10^{-2}$ | $-2.06 \times 10^{-4}$ |
| CLSM | BPAE (F-actin) | $1.48 \times 10^{-2}$ | $-0.65 \times 10^{-4}$ |
| CLSM | BPAE (Mito) | $2.01 \times 10^{-2}$ | $-1.43 \times 10^{-4}$ |
| CLSM | Zebrafish | $9.11 \times 10^{-2}$ | $-2.62 \times 10^{-4}$ |
| CLSM | Mouse Brain | $1.99 \times 10^{-2}$ | $-0.68 \times 10^{-4}$ |
| TPEF | BPAE (Nuclei) | $3.41 \times 10^{-2}$ | $-7.01 \times 10^{-4}$ |
| TPEF | BPAE (F-actin) | $2.66 \times 10^{-2}$ | $-0.88 \times 10^{-4}$ |
| TPEF | BPAE (Mito) | $4.15 \times 10^{-2}$ | $-2.10 \times 10^{-4}$ |
| TPEF | Mouse Brain | $3.36 \times 10^{-2}$ | $-5.67 \times 10^{-4}$ |
| WF | BPAE (Nuclei) | $7.88 \times 10^{-4}$ | $2.03 \times 10^{-4}$ |
| WF | BPAE (F-actin) | $19.15 \times 10^{-4}$ | $2.36 \times 10^{-4}$ |
| WF | BPAE (Mito) | $9.59 \times 10^{-4}$ | $2.53 \times 10^{-4}$ |

From Table 1, the estimated Gaussian noise variance (*b*) for the confocal and two-photon microscope is negative, which is due to an offset in the noisy image. More details can be found in [40] section II (A). For these cases, we set the estimated Gaussian variance to zero. From Table 1, we also observe that the estimated Poisson noise variance (*a*) in confocal and two-photon microscopy is dominating over the Gaussian noise variance (*a ≫ b*). In the case of widefield microscopy, both noise parameters exist (*a, b >* 0), which indicates the mixture of Poisson-Gaussian noise. Also, the Gaussian variance (*b*) is in the range of ∼ 2e-4, whereas the Poisson variance (*a*) varies depending on the signal intensity. This behavior is observed in the widefield microscope, where the *a* value of BPAE (F-actin) is much larger when compared to the *a* value of BPAE (Mitochondria) and BPAE (Nuclei). The estimation of these noise parameters, i.e., the Poisson variance (*a*) and the Gaussian variance (*b*), shows a clear map between different noises in each microscope modality. Hence, we broadly divide the microscopy modalities into two categories: first, confocal and two-photon microscopy with Poisson-dominated noise; second, widefield microscopy with MPG noise. For each class of noise type, we show the trained model performance on a single noisy image and the statistics on the complete test-mix dataset.

In this work, we first select a single test image with Poisson-dominated noise, which is captured by a confocal microscope, and another test image with MPG noise, which is captured by a widefield microscope. Here these images are considered as specific images that do not reflect the complete test dataset.

Figure 4 illustrates a zebrafish embryos image with Poisson-dominated noise [EGFP labeled Tg (sox10:megfp) zebrafish at two days post fertilization] captured with a commercial confocal microscope (More details are provided in the Supplementary Note S1). Similarly, Figure 5 shows an image of a BPAE cell (slide #1, F36924 that contains Nuclei, F-actin, and Mitochondria) with MPG noise captured by a commercial widefield microscope. In these figures, the target intensity images were generated by taking the average of 50 noisy images in the same FOV. We observe that our ML-based Noise2Noise plugin denoising method reduces both Poisson-dominated and MPG noise significantly by providing a PSNR improvement of 10.16 dB and 10.5 dB in zebrafish and BPAE cell, respectively.

**Figure 4:**
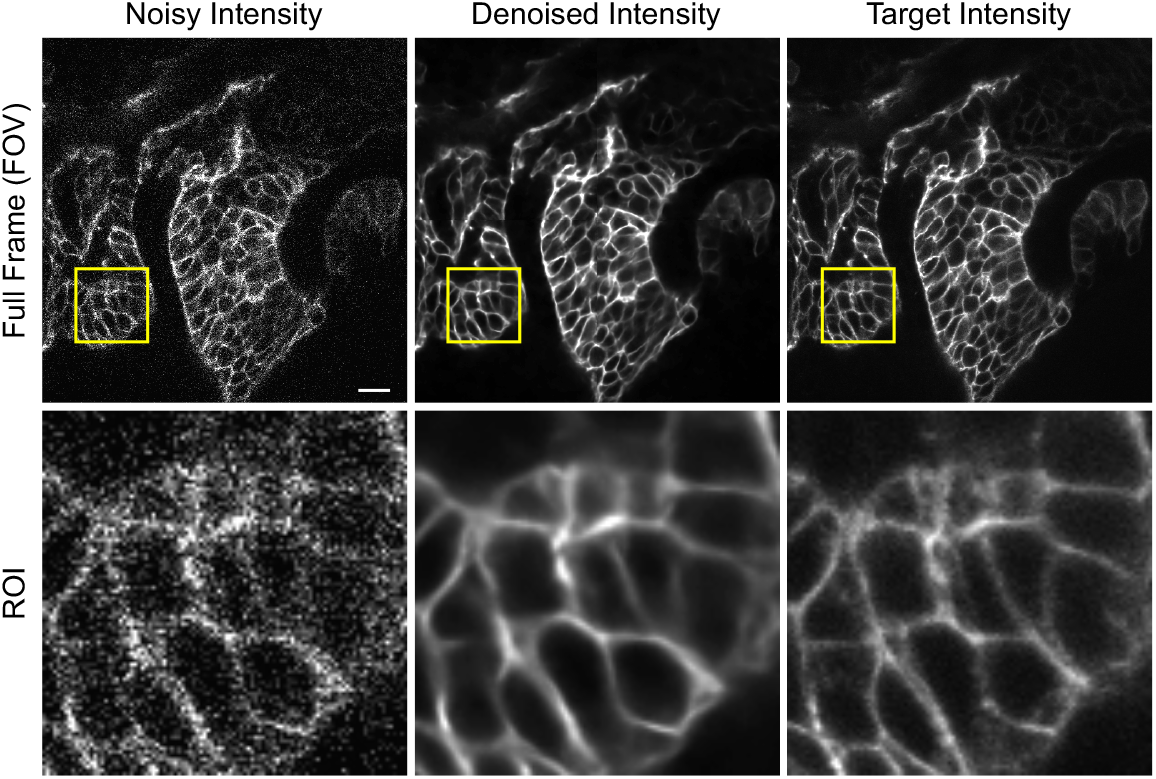
Image denoising results using the Noise2Noise plugin on a fixed zebrafish embryos [EGFP labeled Tg(sox10:megfp) zebrafish at two days post fertilization] captured with confocal microscopy (pixel dwell time of 2 *µ*s and the pixel width of 300 nm, with 10% of power and PMT gain of 140 (model number: Hamamatsu, PMT R10699), excitation and emission wavelengths are 488 nm and 509 nm respectively). The top row indicates the full-frame (of size 512 × 512) of noisy input, denoised output, target, and the bottom row indicates the region of interest (ROI: marked in the yellow square of size 100 × 100) from the respective top row images.

**Figure 5:**
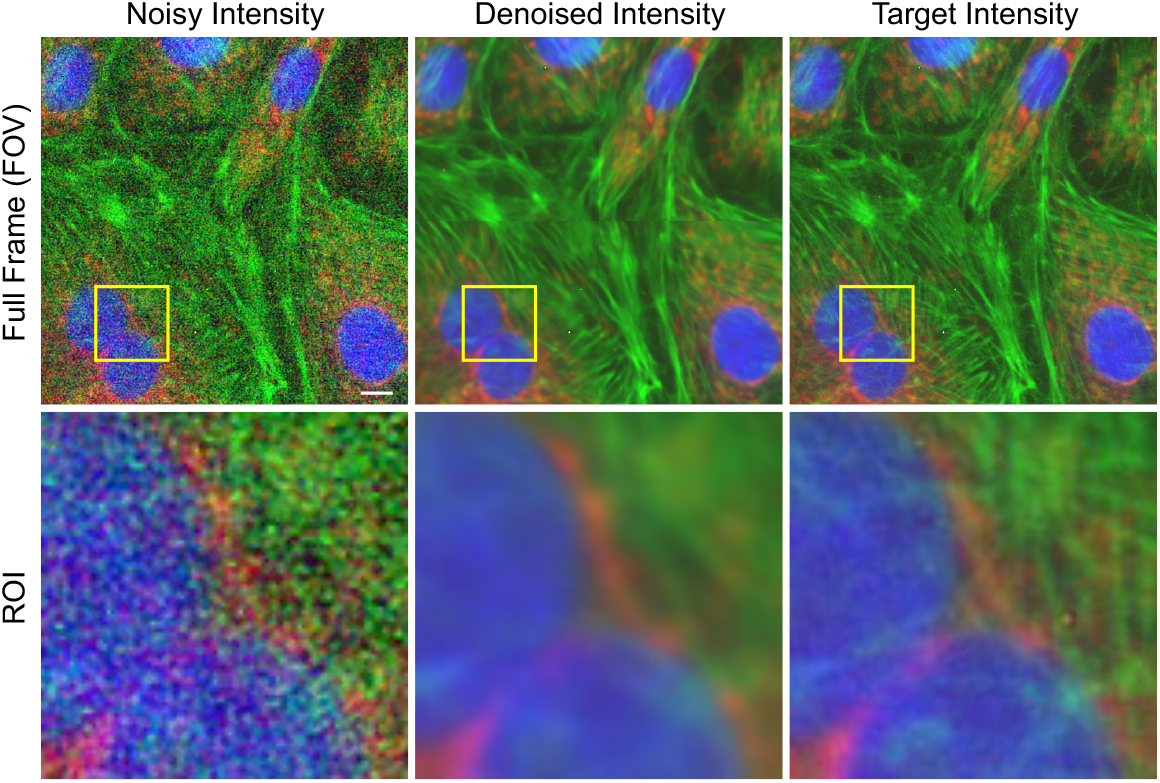
Image denoising results using the Noise2Noise plugin on the BPAE cell [labeled with MitoTracker Red CMXRos (mitochondria), Alexa Fluor 488 phalloidin (F-actin), and DAPI (nuclei) with excitation/emission wavelengths are 561/599, 488/512 and 405/461, respectively; Invitrogen FluoCells prepared slide#1: F36924] captured with the widefield microscopy (exposure time of 200 ms, frame rate of 5 Hz, high definition color camera and controller (DS-Fi1 and DS-U2) with CCD gain of 46 and the pixel width is 170 nm). The top row indicates the full-frame (of size 512 × 512) of noisy input, denoised output, target, and the bottom row indicates the region of interest (ROI: marked in the yellow square of size 100 × 100) from the respective top row images. We imaged three times for the same FOV, each time with a different filter block (DAPI for nuclei, FITC for F-actin, TRITC for mitochondria), to acquire the multi-channel (color) fluorescence image of the cells.

To quantitatively evaluate whether the denoised images contain similar image features as the clean image, we calculate the structural similarity index measure (SSIM) between the two. The SSIM compares luminance, brightness, and contrast values as a function of position [41]. SSIM measures the similarity between two images on a scale of 0 to 1, with 1 being perfect fidelity. In addition, the SSIM can be correlated to the PSNR, and more details are provided in [42]. From the images in Figure 5, the denoised images appear visually similar to the respective target images and with denoised images SSIM values of zebrafish and BPAE samples are 0.84 and 0.83, respectively.

Second, the PSNR distribution of the raw images from the test-mix data and the corresponding denoising results using our ML models are provided in Figure 6. Figure 6 shows the trained Noise2Noise plugin ML model (with an ILR of 5e-4) image denoising performance with complete statistics on the test-mix dataset raw images (48) from our FMD dataset. Also, for image denoising, we train other ML model (requires a clean target), which is a DnCNN ML model that can estimate the residual (only noise) for a given noisy input image. Here, the denoised image is obtained by subtracting the estimated residual from the noisy input data. Figure 6 also shows complete statistics on the test-mix data using the DnCNN architecture ML model with the same ILR of 5e-4. The ML model with Noise2Noise plugin architecture has a better PSNR when compared to the ML model trained with the DnCNN architecture. Hence, the Noise2Noise plugin model demonstrates superior denoising performance compared to the other ML model, and it does not require the clean target image during the training phase. From the raw data distribution, it is evident that there exist two distinct groups of data corresponding to Poisson-dominated noise and MPG noise. The average PSNR of the raw images (including Poisson-dominated noise and MPG noise images) and the corresponding denoised images are 27.22 dB and 35.40 dB, respectively. From Figure 3, the PSNR improvement using the Noise2Noise plugin method is comparable to that of an average PSNR of 8 noisy raw images in the same FOV (with PSNR of 36.02 dB) and thereby reducing the acquisition time approximately by eight fold.

**Figure 6:**
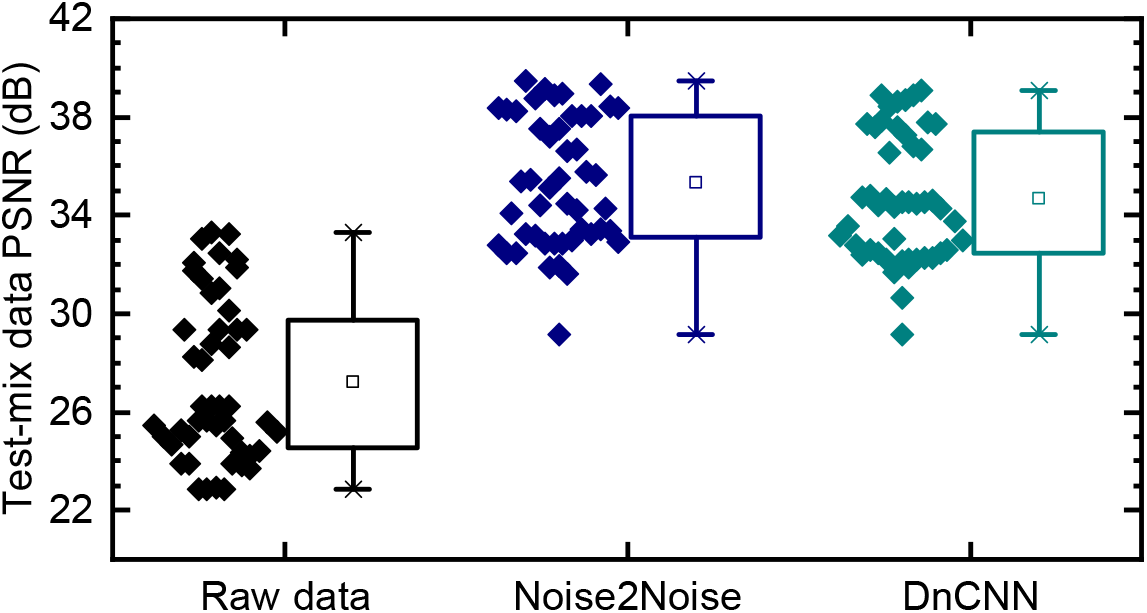
Test-mix dataset (which contains 48 images of the MPG noise samples from different microscopes in the FMD dataset) raw images PSNR box-plot to compare Noise2Noise plugin and DnCNN plugin ML models image denoising performance at a fixed Initial learning rate (ILR: 5e-4). Box-plot contains min, max, 25^th^, and 75^th^ percentile values respectively. Here the square box indicates the mean value. A scattered plot (left to box plot) indicates the true PSNR values used for the statistics.

Finally, we compare the performance of our ML based image denoising methods with the existing image denoising methods. This PSNR improvement is more substantial than existing image denoising methods like PURE-LET, VST+NLM, and VST+BM3D, as shown in Figure 7. Figure 7 shows the existing denoising methods performed on images presented with either Poisson or Gaussian noise. In contrast, our ML based image denoising methods enable superior denoising performance regardless of the noise distribution and the microscopy modality. The PURE-LET, VST+NLM and VST+BM3D image denoising methods perform well only for the images with Poisson-dominated noise, while the NLM and BM3D image denoising methods perform well only for the images with Gaussian noise. In contrast, the ML based image denoising methods (Noise2Noise plugin and DnCNN plugin) perform well for images with both Poisson-dominated noise and MPG noise.

**Figure 7:**
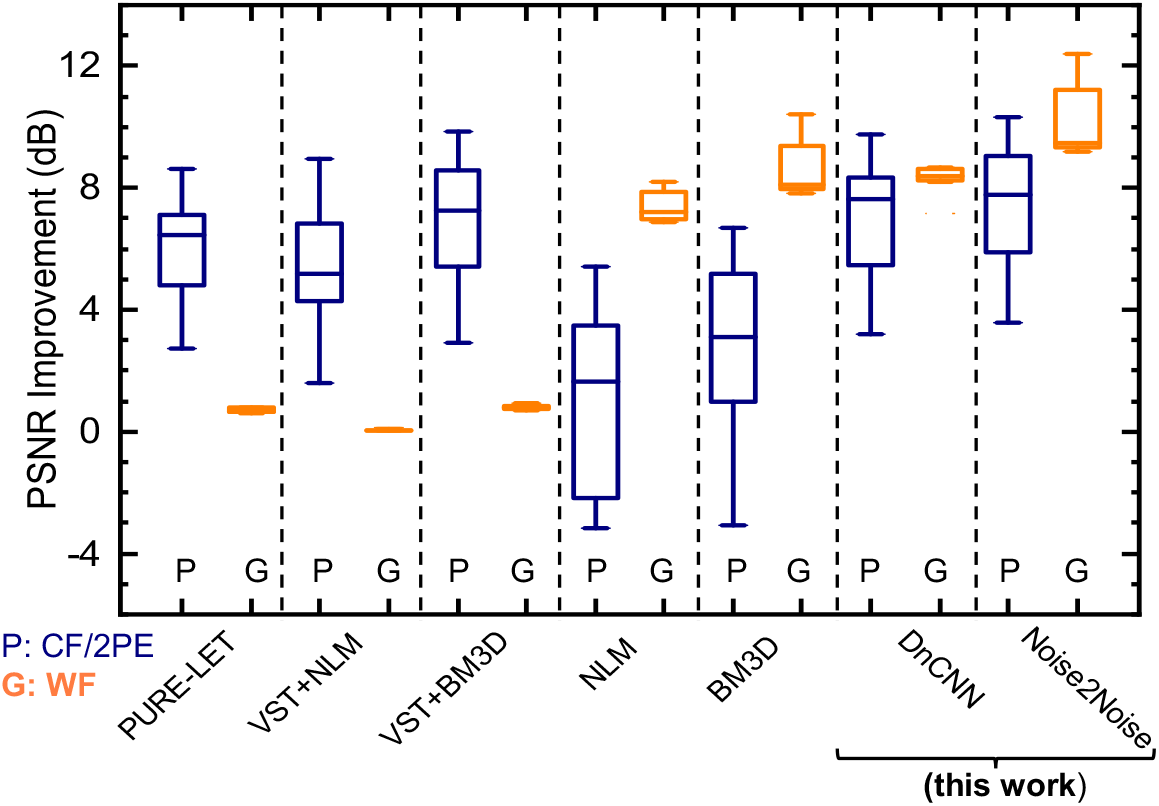
PSNR improvement (denoised image PSNR - raw image PSNR) using different image denoising methods on the raw data with Poisson and MPG noise statistics independently. Here PURE LET, VST+NLM, VST+BM3D, NLM and BM3D are existing image methods and DnCNN plugin and Noise2Noise plugin are our demonstrated supervised trained CNN methods. Blue and Orange color statistics indicate the box plot results of the PSNR improvement for the Poisson-dominated and MPG noises, respectively. Supervised CNN based methods outperform compared to existing image methods independent of noise characteristics.

Our Noise2Noise plugin ML model image denoising method shows significant improvements over existing methods and can provide better results for most of the fluorescence microscopy image follows the MPG noise distribution. To increase the impact and integrate it into existing workflows, we package our method as an ImageJ plugin, a popular image analysis tool used by biologists. Our developed ImageJ plugin has two advantages. First, the plugin can perform image denoising independent of microscopy modality since the ML model was trained with MPG noise images with superior performance compared to existing methods. Second, the plugin can denoise “real-time” (e.g., with an image size of 256 × 256 in 80 ms using GPU) for fluorescence microscopy image with MPG noise. Our plugin denoising time also scales accordingly if the noisy image dimensions are different. More details about the ImageJ image denoising plugin are provided in the Supplementary Note S5. In addition, Supplementary Note S3 shows the qualitative and quantitative comparison of the existing image denoising ImageJ plugins (existing and our ML-based image denoising methods) available with the trained Noise2Noise plugin ML model on a test image from the FMD dataset. The image denoising time (for an image of 256 × 256) using our Noise2Noise plugin on a CPU (Intel Xeon E5-2680 CPU) and on a GPU (Nvidia GeForce GTX 1080 Ti) are 960 ms and 80 ms, respectively. Therefore, the image denoising speed is faster by 12-fold when using GPU compared to CPU. Overall, the ImageJ plugin can be used to denoise fluorescence images that are obtained with low laser power or at a fast acquisition rate. The source code for image denoising including ML models training and testing in Keras and creation of our ImageJ plugin are provided^2^.

## 4 Model Validation

Model validation outside of the training FMD dataset images and noise distributions is crucial to demonstrate the method’s reliability and utility. We validate the Noise2Noise plugin performance (a) on the W2S fluorescence microscopy dataset to compare performance to other analytic and ML-based image denoising methods outside of the FMD training dataset, (b) on the W2S dataset to evaluate denoising performance on samples outside of the training dataset the PSNR distribution and on a flavin-based autofluorescent bacteria images to evaluate performance outside of the training dataset contrast distribution, and (c) discuss the risk and limitations of applying the trained CNN model to higher-dimensional data.

### 4.1 Denoising performance on the W2S dataset

To validate the trained Noise2Noise plugin ML model on images that differ from the FMD dataset, we evaluate denoising plugin performance on 360 widefield fluorescence microscopy images of varying PSNR from the W2S dataset [27]. We also investigate whether the FMD trained ML plugin model applied to the W2S dataset introduces performance degradation or artifacts compared to three of the best performing self-supervised (no external training) ML methods: N2V [20], VST+BM3D [13] and BM3D [9].

The average PSNR improvement and computation time for the different methods applied to the W2S dataset is given in Table 2. For Noise2Noise plugin, the trained ImageJ plugin presented here was used; for the VST+BM3D method (in sequential steps: apply VST transform [13], use the BM3D method [9] and apply inverse VST transformation [13]) for image denoising a and BM3D method [9] MATLAB code was used and for the N2V method an ImageJ plugin was used [43] with two sets of training parameters to explore performance versus computation time trade-offs (“fast” with 10 epochs and 200 steps, “default with 300 epochs and 200 steps). In summary, the supervised Noise2Noise plugin that was trained on the FMD dataset yields higher PSNR improvements compared to self-supervised ML methods. Additionally, the Noise2Noise plugin method only requires convolutions to perform denoising inference, while self-supervised require re-training (or self-training) on every image. The resulting denoising computation time is therefore significantly shorter for the pre-trained Noise2Noise plugin when computed using either a CPU (Intel Xeon E5-2680 CPU) or GPU (Nvidia GeForce GTX 1080 Ti). While the Noise2Noise plugin, VST+BM3D, and BM3D image denoising methods were performed on the entire dataset to find the mean PSNR improvement over the entire W2S dataset, the N2V method requires a long training time per-image, therefore estimates of the mean improvement (and standard error of the estimate) are presented.

**Table 2:** Average PSNR improvement (ΔPSNR= denoised image PSNR - input image PSNR) and single-frame processing computation time (Intel Xeon E5-2680 CPU and Nvidia GeForce GTX 1080 Ti GPU) of Noise2Noise plugin, VST+BM3D, BM3D and Noise2Void image denoising methods when applied to the 360 wide-field fluorescence microscopy images in the W2S dataset. *Estimated mean (and standard error of the mean) from 10 random noisy images from the W2S dataset.

| Method | Training | ΔPSNR (dB) | Time/frame (s)<br>CPU/GPU |
| --- | --- | --- | --- |
| Noise2Noise plugin<br>(this work) | Supervised, FMD | 8.25 | 3.07/0.08 |
| VST+BM3D | Self-supervised | 4.37 | 5.67/None |
| BM3D | Self-supervised | 4.74 | 4.65/None |
| Noise2Void<br>(fast settings) | Self-supervised | 4.6 (0.7)* | 15600/400 |
| Noise2Void<br>(default settings) | Self-supervised | 4.8 (0.7)* | 478800/12000 |

A qualitative comparison of denoising performance is given in Figure 8 for a representative image from the W2S dataset. Here the target image is an average of 400 noisy images in the same FOV and is provided by the W2S dataset. Additional examples are available in Supplementary Note S4 for single-channel and multi-channel test images from the W2S dataset. From these examples, the images qualitatively exhibit lower noise and faithfully represent the target image.

**Figure 8:**
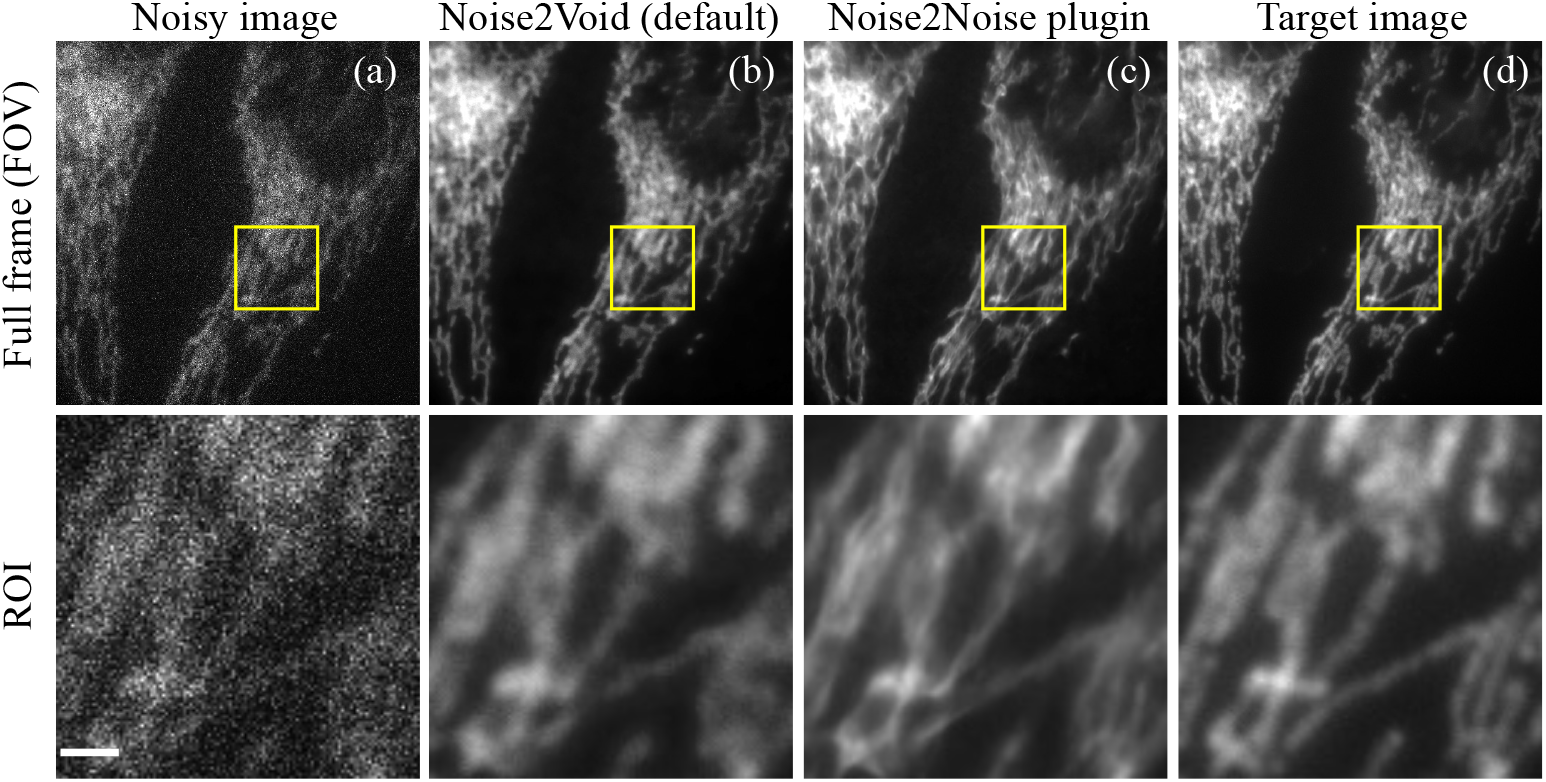
Comparison of the ML-based image denoising plugins in ImageJ. (a) Noisy image, (b) denoised by Noise2Void ImageJ plugin (default training), (c) denoised by the Noise2Noise plugin, and (d) ground truth image of a test mage from the W2S dataset. The top row indicates the full-frame (of size 512 × 512) and the bottom row indicates the region of interest (ROI: marked in the yellow square of size 120 × 120) from the respective top row images. Scale bar: 10 *µ*m.

### 4.2 Out-of-distribution noise and contrast

Machine learning models are optimized for inference data distributions that are similar to the training data distribution. The FMD training dataset is comprised of images with a PSNR of the range of 22-40 dB with contrast ratios (defined as (*I*_*max*_ − *I*_*min*_)*/*(*I*_*max*_ + *I*_*min*_)) of approximately 0.94. To evaluate model performance outside of the training dataset distribution, we validate the trained Noise2Noise plugin on images of varying PSNR and contrast in this section.

The FMD training dataset includes MPG noise from the FMD dataset with PSNR in the range of 22-40 dB; therefore, the model is expected to perform best when for input images with similar noise levels and input images with lower PSNR outside of the training dataset range can expect lower levels of improvement or artifacts [44]. The trained Noise2Noise plugin model presented here, however, is designed to properly denoise images with out-of-distribution noise using “bias-free” (i.e., no bias term) in the convolutional layers [44]. By not including a bias term in the convolutional layers or batch-norm regularization layers, the Noise2Noise plugin provides PSNR improvements when applied to low-SNR images. In this section, we evaluate denoising performance on the W2S dataset for images with out-of-distribution PSNR.

To validate that the bias-free Noise2Noise plugin model can appropriately denoise low PSNR out-of-distribution images, we evaluated denoising performance on such data using the method described in [44]: Gaussian noise is synthetically added to images from the W2S dataset to yield images with PSNR on the range from 5-40 dB, and denoising performance can thereby evaluated as a function of input PSNR. Results are shown in Figure 9 (dark blue symbols and line) show significant PSNR improvements (¿ 9.6 dB) for the W2S test data outside of the trained PSNR distribution (¡ 22 dB). For images with the trained noise distribution range (input PSNR of 22-40 dB, light blue shading), the average PSNR improvement is ≈7.06 dB. PSNR improvement decreases as the input PSNR value increases within the trained noise distribution, and this behavior is consistent with the results shown in Figure 3.

**Figure 9:**
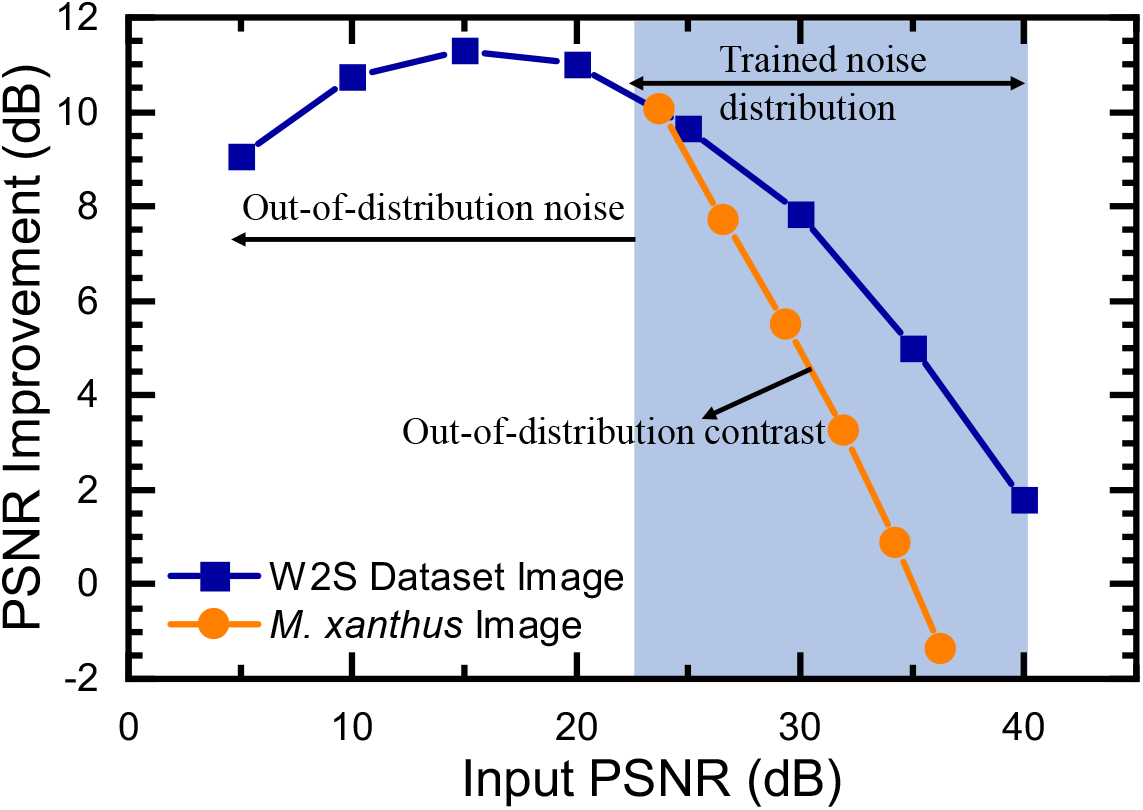
PSNR improvement using the Noise2Noise plugin on test data with acquired PSNR (blue) and contrast (orange) that are out-of-distribution relative to the PSNR and contrast of the FMD training dataset. Out-of-distribution low PSNR images were obtained by adding synthetic noise to the W2S dataset. Out-of-distribution contrast images are taken from the *M. xanthus* autofluorescence dataset and averaged (1, 2, 4, 8, 16, and 32 frames with the same FOV) to obtain images with varying PSNR.

To validate denoising plugin performance for images with out-of-distribution contrast, we evaluate denoising performance for low contrast images (for illustration, defined as (*I*_*max*_ *− I*_*min*_)*/*(*I*_*max*_ + *I*_*min*_)). This metric defines contrast on range from 0 to 1, where 1 is ideal high contrast. The FMD dataset and W2S datasets are both high-contrast datasets with average contrast measures of 0.94 and 0.95, respectively. To evaluate denoising performance for low-contrast fluorescence microscopy images, we prepared an experimental dataset of 600 images (time lapse data) of *Myxococcus xanthus* bacteria auto-fluorescence using widefield microscopy. This sample and imaging method was chosen to produce experimental results with low average contrast (0.52), far outside of the training distribution. Sample preparation and microscope configuration is described in [45, 46]: unlabeled bacteria are illuminated with 458 nm laser to excite flavins in the bacteria, resulting in autofluorescence. The microscope is configured as an widefield epi-fluorescence microscopy using a single dichroic mirror as both excitation and emission filter, resulting in the excitation light leakage and/or back scattering being detected by the imaging EM-CCD. The resulting contrast is therefore relatively low due to the low emission intensity from unlabeled autofluorescence and the weak background suppression from using a single excitation blocking filter. A representative result is shown in Figure 10 (a), (b) and (c), depicting the out-of-distribution noisy *M. xanthus* image, denoised using the Noise2Noise plugin ML model, target images respectively. The target image is defined as the average of 600 noisy images in the same FOV. In this example, applying the Noise2Noise plugin on a single noisy frame from this dataset increases PSNR from 23.72 dB to 33.78 dB.

**Figure 10:**
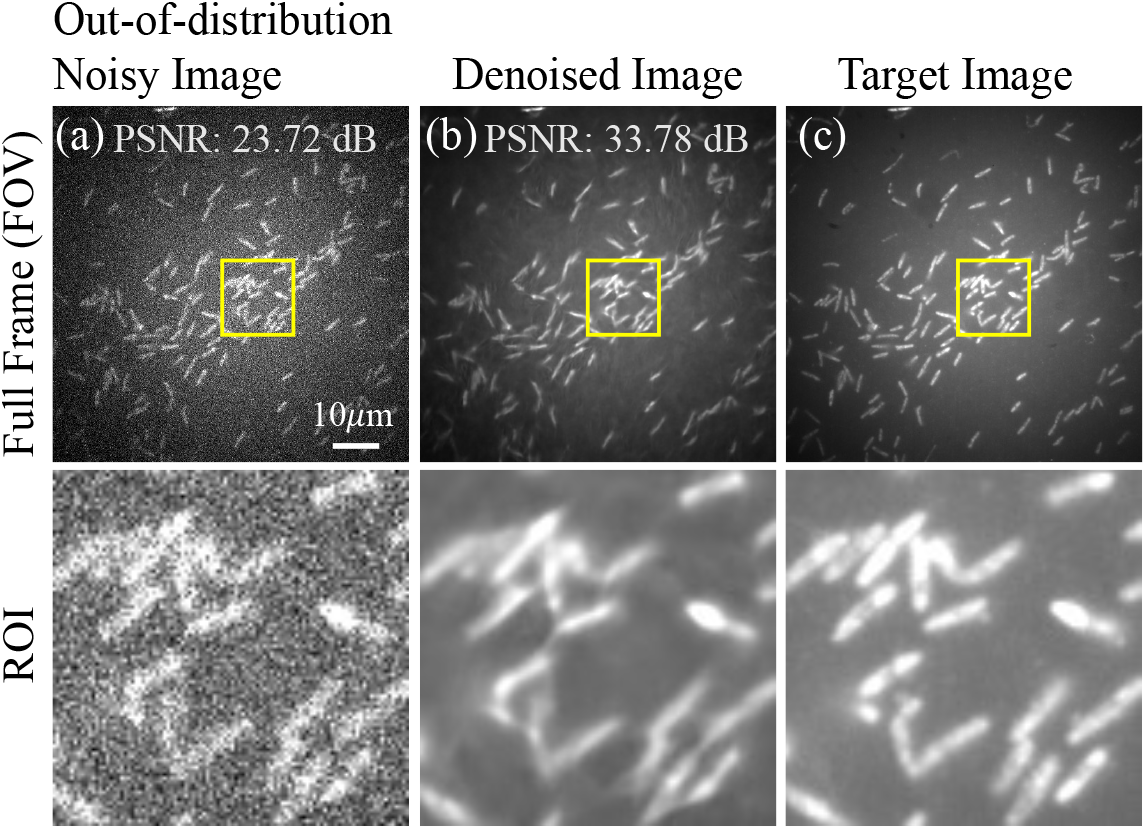
Noise2Noise plugin denoising on *M. xanthus* autofluorescence images captured using a widefield fluorescence microscope. Input noisy image (a), the image denoised using the Noise2Noise plugin (b), and the target image (c) are represented by columns. The full frame FOV is 512 × 512 pixels; the selected ROI (yellow box) is 150 × 150 pixels. Excitation wavelength 458 nm; emission wavelength 500-580 nm; sample excitation power 0.1 mW; integration time 100 ms; pixel size 150nm.

Denoising plugin performance for these low-contrast images can be evaluated as a function of input image PSNR by averaging sequential frames prior to running the denoising plugin. The resulting denoising improvement is given by Figure 9 (orange symbols). The Noise2Noise plugin achieves high-performance denoising on single-frame, low-PSNR microscopy images even when the contrast is far out-of-distribution. While both low-contrast and high-contrast images show decreased denoising performance as input PSNR increases (Figure 3 and Figure 9), denoising improvement decrease at a faster rate for low-contrast images. This defines the out-of-distribution limit for the trained Noise2Noise plugin CNN: exceedingly low-contrast yet high-PSNR input images may have artifacts introduced in to the image.

In summary, the Noise2Noise plugin demonstrates high-performance denoising on out-of-distribution low-PSNR and low-contrast ratio images taken from datasets external to the training dataset. By using bias-free convolutional layers, denoising on low-PSNR images is enabled. The trained CNN performs well on low-PSNR and low-contrast images outside of the dataset, however the CNN introduces artifacts that degrade images in the *M. xanthus* dataset when contrast is low (¡0.6) and PSNR ≳ 36 dB. However, such high-PSNR images would likely not be candidates for denoising, and therefore applying the Noise2Noise plugin is reasonable for practical experimental fluorescence microscopy PSNR and contrast. The Noise2Noise plugin shows exceptional performance (¿8 dB) for single-frame, low PSNR images (¡25 dB) with typical experimental contrast (at least ¿0.5).

### 4.3 Out-of-distribution structures

While the machine learning model was trained on the FMD dataset (containing fluorescent-labeled BPAE cells and intravital genetically encoded fluorescent mouse brain and zebrafish samples), the previous section demonstrated that the model can perform accurate denoising on out-of-distribution *M. xanthus* structures (cellular features that do not appear in the training set images). To further evaluate performance on out-of-distribution structure, the denoising plugin performance was evaluated on the 3D RCAN dataset[33], consisting of fixed U2OS cells (cultivated from the human osteosarcoma bone tissue) transfected with mEmerald-Tomm20 labeling the outer mitochondrial membrane. The organelles are fluorescence stained with Alexa Fluor 488 Phalloidin (actin); ERmoxGFP (endoplasmic reticulum, ER); GalT-GFP (golgi); LAMP1-mApple (lysosomes); and Mouse-*α*-Tubulin primary antibody, Donkey-*α*-Mouse Biotin secondary antibody, and Alexa Fluor 488 Streptavidin (microtubules). In addition to a variety of out-of-distribution structures, the images were acquired using an out-of-distribution imaging modality: “instant structured illumination microscopy” (iSIM) [47]. These structures are chosen due to the diversity of features across length scales and are used for validation in other machine learning models [5]. Hence the 3D RCAN dataset samples were chosen to represent the out-of-distribution structures. To evaluate plugin performance with varying SNR in the RCAN dataset, synthetic Gaussian noise was added to the high-SNR images in the dataset.

Next, the Noise2Noise plugin is applied to the noisy images, and the quantitative improvements in PSNR and SSIM are computed. To account for differing image intensities (dynamic range of pixels values) from the acquired low-SNR and high-SNR samples, normalization is required before calculating the quantitative results. Images are percentile normalized using the method described in [17] and normalized images are denoted as 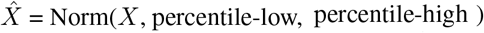 where *X* is the noisy image, second argument “percentile-low” indicates the lowest value of *X* (calculated as percentile(*X*, percentile-low)) and third argument “percentile-high” indicates the highest value of *X* (calculated as percentile(*X*, percentile-high). For noisy (low-SNR)/denoised images percentile-low and percentile-high are set to 1 and 99, respectively. Similarly, target (high-SNR) images percentile-low and percentile-high are set to 0.1 and 99.9, respectively.

**Figure 11:**
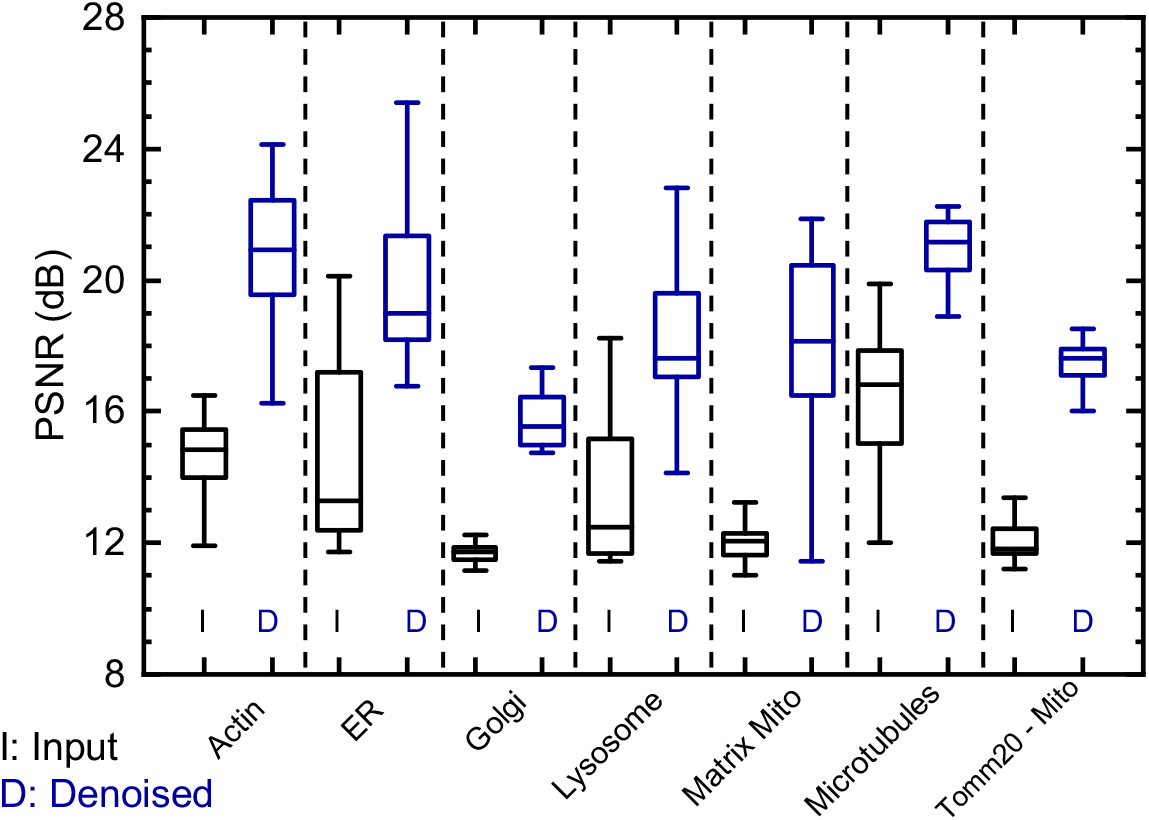
PSNR distribution of the noisy and denoised images taken from the out-of-distribution samples from the 3D RCAN dataset [33]. Gray and Blue color statistics indicate the box plot results of the input and denoised PSNR, respectively.

Table 3 provides a summary of average PSNR and SSIM improvement after denoising on the 3D RCAN dataset [33]. Average PSNR improvements for each class of structures range from 3.83 to 6.39 dB, and each of the 207 images of the dataset yielded a positive increase in PSNR (ΔPSNR ¿ 0) and SSIM (ΔSSIM ¿ 0).

**Table 3:** Average PSNR improvement (ΔPSNR = denoised image PSNR - input image PSNR) and average SSIM improvement (ΔSSIM = denoised image SSIM - input image SSIM) of the Noise2Noise plugin when applied to the out-of-distribution samples of fluorescence microscopy images from the 3D RCAN dataset. Population standard deviations of the improvements are given in parenthesis.

| Sample | Num. Samples | ΔPSNR (dB) | ΔSSIM |
| --- | --- | --- | --- |
| Actin | 21 | 6.39 (2.10) | 0.59 (0.06) |
| ER | 24 | 4.91 (1.35) | 0.32 (0.09) |
| Golgi | 34 | 3.83 (0.84) | 0.21 (0.05) |
| Lysosome | 35 | 4.99 (1.84) | 0.33 (0.09) |
| Matrix Protein Mitochondria | 36 | 6.10 (2.20) | 0.33 (0.16) |
| Microtubule | 32 | 4.38 (2.01) | 0.39 (0.11) |
| Tomm20 Mitochondria | 25 | 5.35 (0.79) | 0.33 (0.06) |

From the Table 3, our Noise2Noise plugin shows better image denoising performance on the out-of-distribution structures commonly arise in the biomedical imaging. Qualitative results on the 3D RAN dataset, and additional out-of-distribution datasets (such as fluorescence microscopy samples of BPAE cells, membrane samples [48]), fixed pattern noise dataset captured using an out-of-distribution modality total internal reflection fluorescence (TIRF) microscope and a lattice light-sheet microscopy (LLSM) microscope systems [5] are shown in Supplementary Note S4 and in our GitHub repository under Model validation folder ^3^.

### 4.4 Application specific model validation

The use of pre-trained neural network models bears the risk of introducing artifacts, especially when applied to images that differ from the training data set. While this paper evaluates Noise2Noise plugin performance on some out-of-distribution contrast, SNR, structure, and imaging modality data sets, applying the plugin generally to untested data requires the users to examine performance. The following steps can be applied in order to evaluate performance on new data sources, samples, or microscopy modalities. Application-specific validation follows three steps: image acquisition, applying denoising through ML plugin, and performance verification.

First, obtain several single-channel low-SNR frames of the same fluorescence microscopy image field of view (i.e., at least greater than ten 2D sections at low excitation power or fast data acquisition rates). In addition to each individual frame, an estimate of the clean image is to be obtained by either acquiring the same field of view under long-term exposure or by averaging all the acquired low-SNR frames together. To obtain the average, the collection of individual frames can be combined in Fiji/ImageJ as a “Stack,” then averaged using the “Stack” menu commands. Alternatively, a high SNR estimate can be obtained by performing averaging on the individual frames using a variety of software tools (e.g., Python, MATLAB). The output representing the clean image is then saved to disk.

Second, load individual frames into Fiji/ImageJ and apply the Noise2Noise plugin. The installation of the Noise2Noise plugin and other details are provided in the Supplementary Note S5 and GitHub repository. Save the output denoised image files to disk.

Third, evaluate denoising performance by comparing the output of the denoising plugin to the estimated clean image. To evaluate performance qualitatively, first load both images simultaneously in Fiji/ImageJ while ensuring that both images are displayed with the same “Brightness/Contrast” scale (e.g., in the menu *Image→Adjust→Brightness/Contrast*…). Next, compare the structure of specific features of interest between the two images as well as examining the background (i.e., low-intensity) regions of the image for apparent artifacts. Quantitative verification can be performed by computing and comparing the PSNR and SSIM metrics of the denoised images with respect to the clean images. PSNR and SSIM for the raw images compared to the clean image and the plugin output compared to the clean image can be calculated using ImageJ plugins [49, 50]. The PSNR and SSIM of the output of the denoising plugin is expected to achieve at least ≈3-8 dB improvement in PSNR and a positive increase in SSIM compared to the raw noisy images. Additional examples and detailed explanations of Noise2Noise plugin are provided in the GitHub repository.

### 4.5 Out-of-model data: higher dimensionality data

Modern microscopy techniques commonly acquire data in four or more dimensions (three spatial, one temporal, and possibly more channels) and represent the data as 2D sections per channel at discrete acquisition times. It is therefore important to consider the model in the context of data with dimensionality outside-of-distribution from the training dataset. To address this, the model was trained on 2D sections of 3D spatial data of one to three channels, captured at specific times during dynamic intravital processes (e.g., blood flow and respiration). The noise of each individual channel can be described as MPG noise, and therefore it is appropriate to apply the method to 2D sections of higher dimensional data using the MPG denoising method. Since the trained Noise2Noise plugin model performs denoising solely using information from a single 2D plane of the 3D sample, potential further performance improvements could be achieved by developing and training models using complete high-dimension data (e.g., the entire 3D stack in time [51]). Such an approach relies on significant computational improvements and would effectively amplify signals correlated in 3D space and time, leading to even improved denoising performance.

## 5 Conclusion

In this paper, we have presented an open-source software tool for fluorescence microscopy image denoising in real-time (within tens of milliseconds) to denoise mixed Poisson-Gaussian noise. The new image denoising technique using the trained ML models (Noise2Noise plugin and DnCNN plugin) allows imaging eight times faster than the fundamental speed limit of fluorescence microscopy while maintaining the same PSNR. The system was trained using the experimentally obtained FMD dataset to develop a neural network that effectively removes MPG noise presented in widefield, confocal, and two-photon fluorescence microscopy images across a variety of cell types. We have shown here that the proposed Noise2Noise plugin ML model demonstrates larger PSNR improvements compared to the existing image denoising methods, performs denoising faster, and has been validated on a wide variety of images varying SNR, contrast, structures, and imaging modality.

## Data and code availability

The FMD dataset mentioned in this paper is publicly available in the CurateND: https://curate.nd.edu/show/f4752f78z6t. The code for training the Noise2Noise plugin and DnCNN plugin architectures and underlying the results presented in the paper are publicly available in the GitHub repository: https://github.com/ND-HowardGroup/ Instant-Image-Denoising/. Also, this repository includes the estimation of noise parameters using MATLAB codes, Noise2Noise plugin validation on W2S dataset, Noise2Noise plugin validation on out-of-distribution samples, and plugin source code in java with ImageJ plugins. Additional details of our Noise2Noise plugin denoising validation on the large datasets are available in CurateND: https://curate.nd.edu/show/5h73pv66h5f. Out-of-distribution structure dataset (3D RCAN dataset) used to generate underlying denoising results presented in this paper are available in Ref. [33].

## Acknowledgments

We thank Prof. Joshua D. Shrout, Department of Civil and Environmental Engineering and Earth Sciences, University of Notre Dame, for providing *M. xanthus* cells for our experiments. Yide Zhang’s research was supported by the Berry Family Foundation Graduate Fellowship of Advanced Diagnostics & Therapeutics (AD&T), University of Notre Dame. The authors further acknowledge the Notre Dame’s Center for Research Computing (CRC) for the use of GPUs to train the ML models and the Integrated Imaging Facility (NDIIF) for the use of the Nikon A1R MP confocal microscope and Nikon Eclipse 90i wide-field microscope in NDIIF’s Optical Microscopy Core.

## Footnotes

2 https://github.com/ND-HowardGroup/Instant-Image-Denoising

3 https://github.com/ND-HowardGroup/Instant-Image-Denoising/tree/master/Plugins/Model_validation

