## Supplemental Document for "Real-time Image Denoising of Mixed Poisson-Gaussian Noise in Fluorescence Microscopy Images using ImageJ"

### Realtime image denoising of mixed PoissonGaussian noise in fluorescence microscopy images using ImageJ: Supplemental document

February 8, 2022

#### NOTE S1: Fluorescence microscopy denoising (FMD) dataset

##### Image denoising using machine learning

The image acquisition speed limits image peak signal-to-noise ratio (PSNR) in fluorescence microscopy. There exists a variety of image denoising methods to overcome this fundamental limitation. However, there are few drawbacks with these methods such as requirement of noise statistics (variance information), poor denoising performance and computationally expensive. Recently machine learning (ML) approaches show a dominant performance improvement in imaging-related tasks such as classification or/and segmentation, and also enabling faster computation time [1]. Although these ML models are superior in performance compared to conventional methods, there are a few limitations as follows. First, the ML models are data-driven and require an extensive training data which are relatively difficult to collect, and second, the ML models require an extensive training time along with expensive resources like GPU's. Here, addressing the first fundamental limitation is vital because to generalize the ML model one has to train with various imagining systems (microscopic modalities and different samples). Hence, for training different ML models, one require a large fluorescence microscope image denoising dataset. We address this fundamental problem by providing the fluorescence microscopy denoising (FMD) dataset[1] which contains 12,000 raw noisy images from confocal [2], two-photon [3], and widefield [4] microscopes. In this work, we train the Noise2Noise plugin and DnCNN plugin ML models using the FMD dataset [5]. The collected FMD dataset by our group is also helps to benchmark various image processing techniques on the fluorescence microscopy dataset.

[1] https://curate.nd.edu/show/f4752f78z6t, DOI:10.7274/r0-ed2r-4052

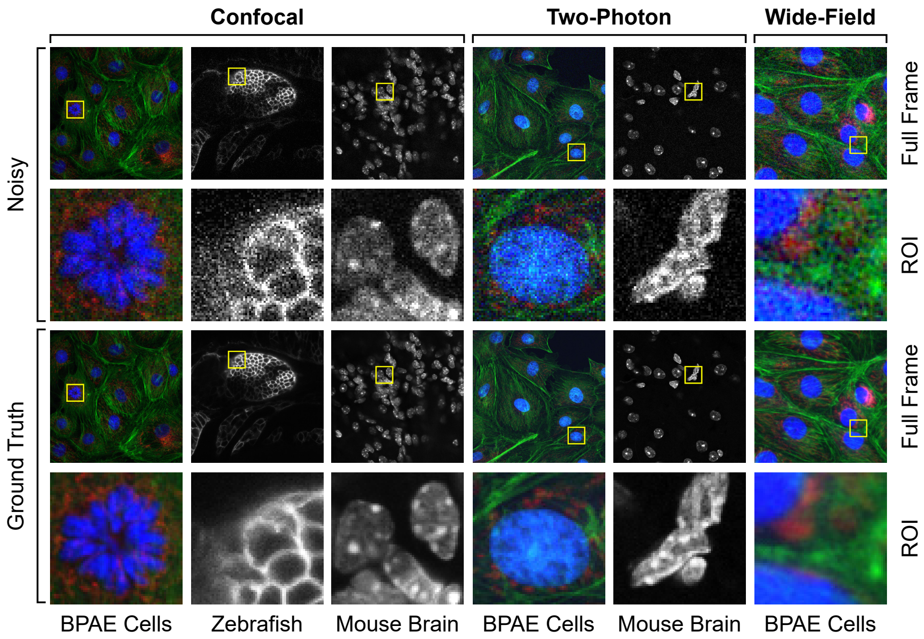


**Figure 1:** Illustration of raw fluorescence microscopy images and their estimated ground truth from the FMD dataset. Shown here are FOVs from different microscopy modalities on real biological samples [6].

#### Data acquisition and Image registration

To create the FMD dataset, we imaged different real biological samples like fixed cells (BPAE cells) and *in vivo* images (mouse brain and zebrafish) using various microscopy modalities such as confocal, two-photon, and widefield microscopes. All animal studies were approved by the university's Institutional Animal Care and Use Committee. Figure 1 shows the illustration of the various microscopy modalities and corresponding images samples (total 12 samples). More details about the acquisition setup are available[2]. Here, each biological sample consist of 20 different field of views (FOVs) and at each FOV we capture 50 noisy images. Therefore, for 12 biological samples, it would result in 12000 raw noisy images. To include images with different PSNR, we perform data augmentation using the raw noisy images, i.e., for each image in a FOV, we consider an average of 2, 4, 8, and 16 consecutive images within a single FOV which eventually results with additional 4 more images. Also, taking an average of $N$ images leads to a high PSNR image as $N$ increases to 16 in an increasing order. This is due to the fact that averaging suppress the noise. Therefore, by this approach of data-augmentation, we increased the FMD dataset to 60,000 images with various PSNR values.

[2] `https://github.com/ND-HowardGroup/Instant-Image-Denoising/tree/master/Image%20acquisition%20setup`

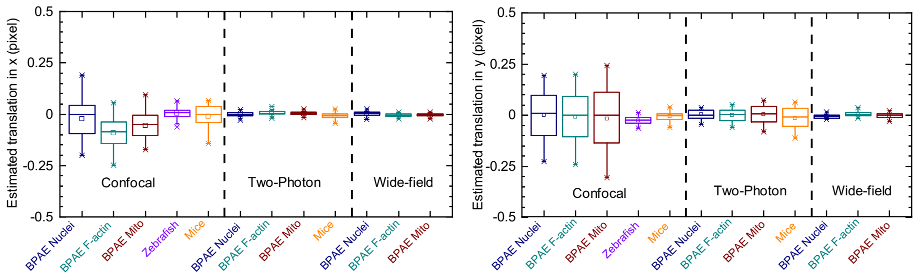


**Figure 2:** Estimated translation in $x-$ and $y-$ axis are both within a half-pixel (0.5), as shown in the left and right images, respectively. The estimation statistics represent data of the complete 20 FOV's of each imaging configuration.

To perform the averaging, one need to show the requirement of image registration on these acquired fluorescence microscopy images. Here, the image registration is required when there is a translation of more than half-pixel in $x-$ axis and $y-$ axis. In this work, we first prove that there is no translation of more than half-pixel in $x-$ and $y-$ axis and hence, the need for the image registration is not required. Figure 2 shows the estimated translation in $x-$ axis and the $y-$ axis along with its statistics. We observe that all imaging modalities have less than half (0.5) pixels shift. This result allow us performing averaging $N$ images within a single FOV and thereby resulting in a high-PSNR compared to an individual noisy image. Also, the clean image in each FOV is obtained by averaging 50 noisy images. The generated clean image is then used for the performance comparison of different ML models. Therefore, after the FMD dataset creation, different ML models are designed to denoise the FMD test dataset. Hence, these trained ML models can be considered as a generalized framework to denoise any fluorescence microscopy image.

### NOTE S2: Training and testing results with ML models

#### Noise2Noise plugin model

In this work, we train a deep convolutions neural network (CNN) based on the Noise2Noise architecture [7] on the FMD dataset. Here, we do not consider the batch-norm between convolutional layers (NBN). We develop the Noise2Noise plugin model using the Keras with TensorFlow as backend [8] with a noisy image and another noisy image in the same field of view (FOV) as the input and target image, respectively. The ML model is trained using Keras, a high-level API of the Tensorflow machine learning package. In the FMD dataset, 95% images are used for the training, whereas the remaining 5% images are used for the testing. During training Adam [9] is the optimizer to update the convolutional neural network kernel coefficients (weights) with 200 epochs (iterations), including the training batch size of 4. Also, the additional settings during the training process are used: first applying weights initialization (drawn from the orthogonal distribution instead of a normal distribution of weights) and second using adaptive learning rate with the Keras One-cycle [10] method.

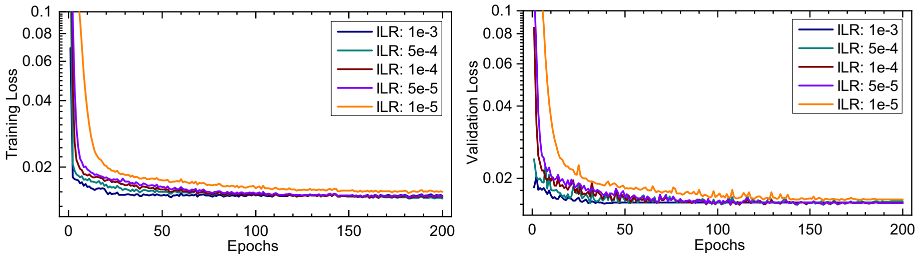


**Figure 3:** Training and validation loss (shown in the left and right plots, respectively) over epochs of the trained Noise2Noise plugin ML architecture with different initial learning rates on the FMD dataset.

Aforementioned, we train the Noise2Noise plugin model with the FMD dataset. Here, we consider the mean square error (MSE) as the loss function between the predicted image and target image. As illustrated in Figure 3, the model was trained with various initial learning rates (ILR: 1e-3, 5e-4, 1e-4, 5e-5 and 1e-5) and we observe that the training is converged within 150 epochs without any over-fit (see the right side of the image that shows the validation loss). To select the best model with an ILR is determined with the high test-data PSNR value. Once the training is completed, we then freeze the CNN model weights and performed the inference on the test-mix data in the FMD dataset. Figure 4 shows the average PSNR of the test-mix raw images for different ILR values along with its statistics. From the statistics, the trained ML model with ILR value of 5e-4 performs better when compared to other ILR values.

| | Average PSNR (dB) on the Test-mix raw-data | | | | |
|---|---|---|---|---|---|
| Initial Learning Rate | 1e-3 | 5e-4 | 1e-4 | 5e-5 | 1e-5 |
| **Noise2Noise-NBN** | 35.07 | **35.35** | 35.32 | 35.18 | 34.20 |
| Noise2Noise-BN | 34.98 | **35.03** | 34.75 | 34.53 | 33.21 |
| DnCNN | 34.57 | **34.67** | 34.59 | 34.29 | - |

**Table 1:** Average PSNR (dB) on test-mix data raw images using the FMD dataset with the trained machine learning models (Noise2Noise-NBN, Noise2Noise-BN, and DnCNN) at initial learning rates (ILR).

Also, we train the Noise2Noise plugin model by enabling the batch-norm (named as “Noise2Noise-BN”) between the convolutional layers, and evaluate its performance across different ILR and summarized in Table 1. From this Table, we observe that if we disable the batch-norm slightly helps in average PSNR on the test-mix data from the FMD dataset across different ILR values.

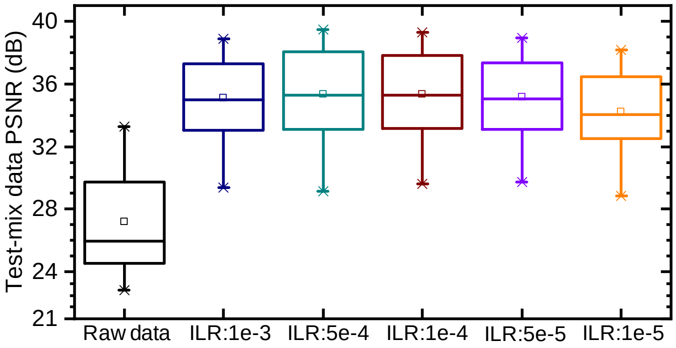


**Figure 4:** Test-mix dataset (from the FMD dataset) raw images PSNR using the Noise2Noise-NBN model with different ILR values. From this box-plot, it is evident that at the trained model with ILR: 5e-4 results in better PSNR compared to other models. Box-plot contains min, max, $25^{\text{th}}$, and $75^{\text{th}}$ percentile values respectively. Here the square box indicates the mean value.

#### DnCNN plugin model

Here, we generated the clean target images (average of 50 noisy images with in the same FOV) from the FMD dataset that can be used for PSNR improvement estimation of the trained ML model. For image denoising, we train other ML model (that requires a clean image), which is denoising using a convolutional neural network (DnCNN [11]) that can estimate the residual (only noise) for a given noisy input image. The architecture of the DnCNN plugin model contains 17 convolutional layers with 64 channels in each layer. Each convolutional layer contains convolution operation, ReLU and followed by the Batch-norm layer. Here, the denoised image is obtained by subtracting the estimated residual from the noisy input. The PSNR performance of the DnCNN plugin model is also provided in Table 1 with different ILR values. From this Table, it is evident that the Noise2Noise plugin ML model performs better compared to the DnCNN plugin ML model at all ILR values. Consolidated results along with the source code are available[3].

### NOTE S3: Comparison of existing image denoising methods in ImageJ

We broadly divided the image denoising methods into two sections, traditional methods and ML-based methods in ImageJ.

[3] `https://github.com/ND-HowardGroup/Instant-Image-Denoising/tree/master/Python`.

#### Traditional image denoising methods

Conventional image denoising methods like NLM, BM3D are explained in the main text. Here we present a list of conventional image denoising plugins in ImageJ. A couple of image denoising plugins in ImageJ perform similarly to the Gaussian denoising method. A few of these methods are listed as: ROF denoising [12,13], Non-local means denoising (NLM) [14,15], Pure Denoise [16], Wavelet denoising [17], and Candle-J denoising [18]. Initial comparison of our pre-trained ML Noise2Noise plugin model with ROF denoising and NLM denoising for the gray channel and color channels, respectively, are provided in our previous paper [19].

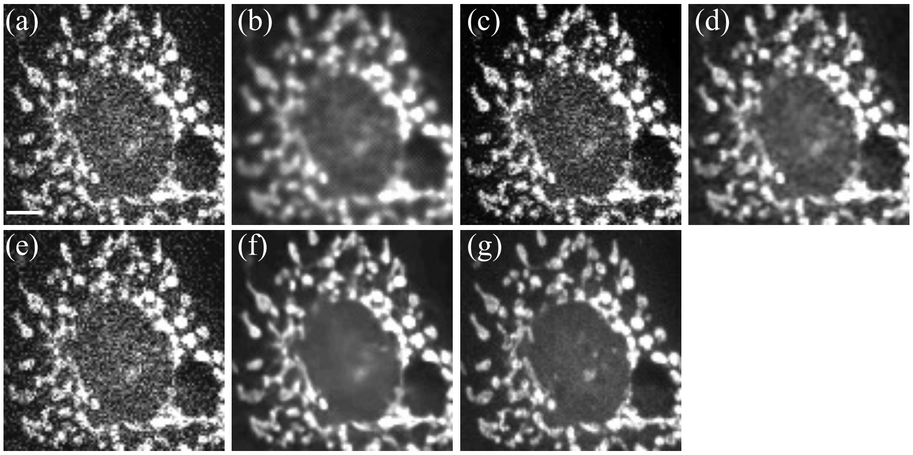


**Figure 5:** Comparison of the traditional image denoising plugins in ImageJ. (a) Noisy image, (b) ROF denoising, (c) NLM denoising, (d) Pure denoising, (e) Wavelet denoising, (f) denoising using the Noise2Noise plugin, and (g) ground truth image of a BPAE sample captured using confocal microscopy. Image dimensions, 100 $\times$ 100 pixels; pixel size, 250 nm; scale bar, 10 $\mu$m.

Figure 5 shows the images denoised by the plugins mentioned above with default settings on a test sample (BPAE sample here) captured using confocal microscopy (red channel). Here the noisy image has a PSNR of 23.29 dB, which is shown in Figure 5(a). ROF denoising results in a PSNR of 28.43 dB, shown in Figure 5(b). Similarly, NLM denoising, Pure denoising, and Wavelet denoising methods result in PSNR values of 24.69 dB, 28.37 dB, and 23.28 dB, which are shown in Figs 5(c), (d) and (e), respectively. Finally, demonstrated the Noise2Noise plugin image denoising method results in a PSNR of 28.80 dB, which is the greatest in performance with computation time in 80 ms (using GPUs) as shown in Figure 5(f). The PSNR comparisons are made with the target image, which is generated by averaging 50 noisy images within the same FOV (Figure 5(g)).

#### ML-based image denoising methods

We also add recently developed ML-based image denoising methods and divide these methods into two categories: those using pre-trained ML models and those using self-supervised ML models. In the first case, image denoising is performed using models pre-trained using a dataset which uses noisy images as inputs and corresponding ground truth images as targets. In the second case, image denoising is performed using the same training dataset and Noise2Void method [20] is an example in this scenario. In this subsection, we compare Noise2Void [20, 21] ML model with the Noise2Noise plugin ML model.

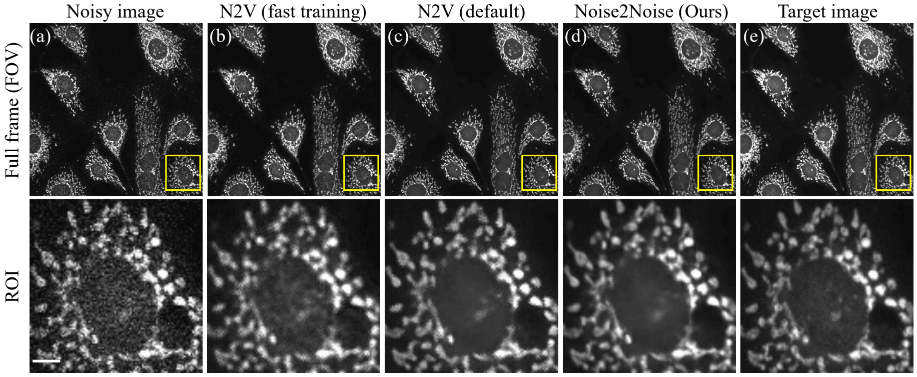


**Figure 6:** Comparison of the Noise2Void ML-based image denoising plugins in ImageJ. (a) Noisy image, (b) Noise2Void (fast training) denoising, (c) Noise2Void (default settings) denoising, (d) Noise2Noise plugin, and (e) ground truth image of a BPAE sample captured using confocal microscopy with 512 $\times$ 512 pixels as image dimension and 250 nm pixel size. The top row shows the full-frame and the bottom row indicates the region of interest (ROI, marked in the yellow square of size 100 $\times$ 100 pixels) from the respective top row images. Scale bar, 10 $\mu$m.

Figure 6 shows the images denoised by the Noise2Void and Noise2Noise plugin (ours) ImageJ plugins as mentioned earlier of a test sample (BPAE sample here) captured using confocal microscopy (red channel). Here the noisy image PSNR is 26.87 dB which is shown in Figure 6(a). Noise2Void is a self-supervised image denoising method. Hence, for a given single image, the method can train and predict on the same image. However, Noise2Void requires training with a certain number of iterations (epochs and steps), which can be divided into two cases: first, fast training with reduced settings of epochs (10 epochs and 20 iterations), and second, default settings (300 epochs and 200 steps) as mentioned in the N2V ImageJ plugin [21]. Figure 6(b) shows the image denoised using the faster training case with a PSNR of 29.81 dB. For the default settings case, the denoised image PSNR is 30.40 dB and is shown in Figure 6(c). The computation time for the two cases are 26 minutes and 8100 minutes (around 5 days and 13 hours), respectively, using a CPU system. On the other hand, Figure 6(d) shows the image denoised using the Noise2Noise plugin with a PSNR of 31.16

dB with computation time of only 3 seconds on the same CPU system. All these PSNR calculations are performed using the target image (by taking an average of 50 noisy images within the same FOV) as reference (Figure 6(e)). Compared to the default settings of Noise2Void, Noise2Noise plugin ML method shows an improvement of 0.8 dB in PSNR. The CNN's training using CPU systems are taking a long time and graphical processing units (GPUs) are required to enable a fast training process. We repeated the same experiment using a GPU system and the training time for the default settings is reduced to 200 minutes (3 hours and 20 minutes) and the PSNR performance is similar the previous CPU training case.

In summary, the Noise2Noise plugin ML model is performing better than the Noise2Void ML model, which is also validated by the authors of Noise2Void using the BSD68 dataset (see **Figure 7** in [21]). Finally, we note that the Noise2Void ML method is a self-supervised method, which is particularly useful when the training dataset is not available. In comparison, our pre-trained ML methods can skip the training phase and directly perform the inference stage for image denoising.

### NOTE S4: Performance on outside of the FMD dataset

This section provides additional examples to demonstrate the generalization and universality of the trained Noise2Noise plugin ML model by denoising images outside of the FMD dataset. Figure 7 shows the performance of the Noise2Noise plugin ML model on a sample from the widefield to structured-illumination microscopy (W2S) dataset. W2S dataset contains noisy images of different PSNR (average 1, 2, 4, 8, 16, 400 images within a single FOV). Figs. 7(a) to (c) show the single channel results (noisy image, denoised image, and target images, respectively) and Figs. 7(d) to (f) show the multi-channel results in the same order. The single-channel and multi-channel denoised images have better quality and closely match with target images. Here the noisy image is the image without average in the FOV, whereas the target image is an average of 400 noisy images in the same FOV. ML-based denoising operation is performed on the raw and noisy image. A small region of interest (ROI) is included in the bottom row for better visualization. For more details about the sample (human cells) and W2S dataset image acquisition details, please refer to Ref. [22].

Another dataset named here onwards as "GigaDB dataset" [23] includes BPAE samples of three different channels such as mitochondria, F-actin, nucleus, and membrane structures. Figure 8 shows the qualitative results on the GigaDB dataset sample images (taken randomly). Quantitative measurements such as PSNR and structural similarity index measure (SSIM) improvement of the denoised image on the complete GigaDB dataset are provided in Table 2. Here the PSNR value is computed using the percentile based normalization method as explained in [24]. The noisy (low-SNR) and ground truth (high-

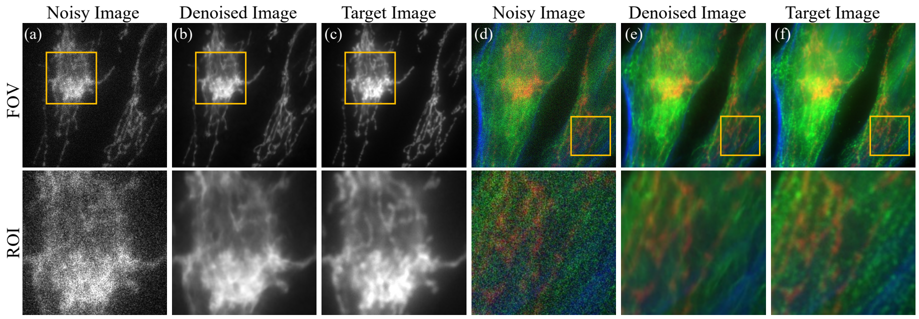


**Figure 7:** Application of the Noise2Noise plugin on a test sample from the W2S dataset [22]. From Left to right: noisy images (a) and (d), denoised images (b) and (e) using Noise2Noise plugin, and target images (c) and (f) of human cells for a single channel and multi-channel images, respectively (the full-FOV of size 512 × 512 pixels). Selected ROI (in the yellow color region) for each image is shown in the bottom row. Scale bar, 10 $\mu$m.

| **Sample** | **Num. Samples** | **ΔPSNR (dB)** | **ΔSSIM** |
|---|---|---|---|
| Nucleus | 104 | 2.34 | 0.26 |
| Membrane | 84 | 1.96 | 0.15 |
| Actin (confocal) | 79 | 3.32 | 0.17 |
| Mito (confocal) | 79 | 5.08 | 0.18 |
| Actin (60x) | 100 | 4.38 | 0.36 |
| Mito (60x) | 100 | 5.60 | 0.39 |

**Table 2:** Average PSNR improvement (ΔPSNR = denoised image PSNR - input image PSNR) and average SSIM improvement (ΔSSIM = denoised image SSIM - input image SSIM) of our Noise2Noise plugin image denoising methods when applied to the samples of fluorescence microscopy images in the GigaDB dataset. Estimated values are the mean of the number of samples column images from the GigaDB dataset [23].

SNR) images are normalized using percentiles (1, 99.5) and (0.1, 99.9) for the low SNR and high SNR images, respectively).

Some more out-of-distribution structures of fluorescence microscopy samples are taken from the 3D RCAN dataset [25] that includes Actin, ER, Golgi, Lysosome, Matrix-mitochondria, Microtubules, Tomm20-mitochondria samples. Here the noisy images are generated by synthetically adding a Gaussian noise on the deconvolution target images from the 3D RCAN dataset. Figure 9 shows the qualitative results on the 3D RCAN dataset sample images (taken randomly). Quantitative measurements of PSNR and SSIM are provided in the main paper. For the complete dataset including noisy and denoised images is provided in the GitHub repository and [4].

[4] https://curate.nd.edu/show/5h73pv66h5f

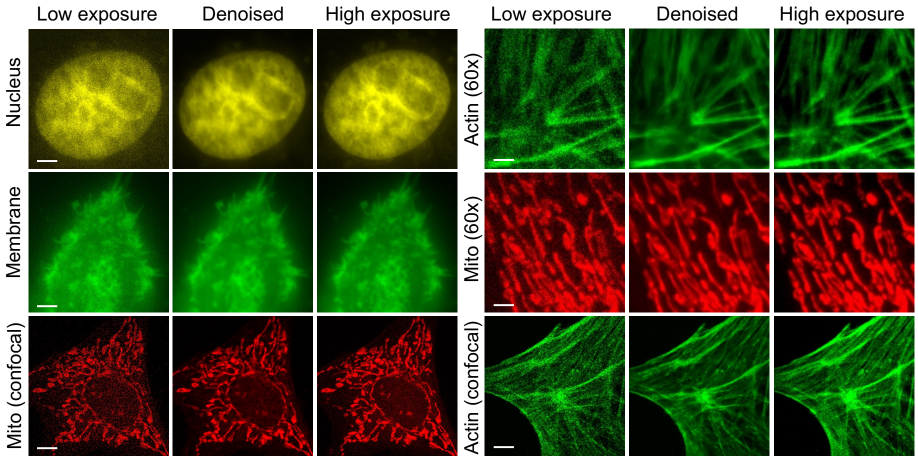


**Figure 8:** Qualitative results of the Noise2Noise plugin on a random test sample from the GigaDB dataset [23]. From Left to right: noisy (low exposure time) images, denoised images using Noise2Noise plugin, and target images (longer exposure time) of BPAE samples (shows nucleus, Actin, and Mitochondria) as single-channel images, respectively. Scale bar, 10 $\mu$m.

In addition to the mixed Poisson-Gaussian (MPG) noise, fluorescence microscopy systems that use CCD or CMOS as detector arrays adds additional noise called “fixed-pattern noise”. Fixed-pattern noise (FPN) is because the detector gain and offset are different at each pixel in the acquired image and can be corrected using calibration (correct the offset then divide by the gain factor). Even our trained models trained with only MPG noise, the Noise2Noise plugin image denoising on the fixed-pattern noise is shown here using out-of-distribution structures such as HeLa cell microtubules taken from the ACsN dataset [26]. Figure 10 shows the qualitative results on the HeLa cell microtubules captured at different integration times (5ms, 10 ms) with FPN using an out-of-distribution modality total internal reflection fluorescence (TIRF) microscopy system.

To provide the quantitative results, one of the HeLa cell microtubules samples is captured using an out-of-distribution modality TIRF microscopy system at two different integration times, with the first one at 10ms and another one at 110 ms. Consider the 10ms integration image as a raw (noisy) image, and the Noise2Noise plugin is used to perform the image denoising on the fixed pattern noise. Figure 11 shows the qualitative results on the HeLa cell microtubules as noisy (10ms), denoised using Noise2Noise plugin and the ground truth (110 ms), respectively. Here the PSNR value is computed using the method explained in [24]. The noisy (low-SNR) and ground truth (high-SNR) images are normalized using percentiles (1, 99.5) and (0.1, 99.9) for the low SNR and high SNR images, respectively). In addition, Figure 12 shows the qualitative results on the adult brine shrimp sample with FPN captured using an out-of-distribution

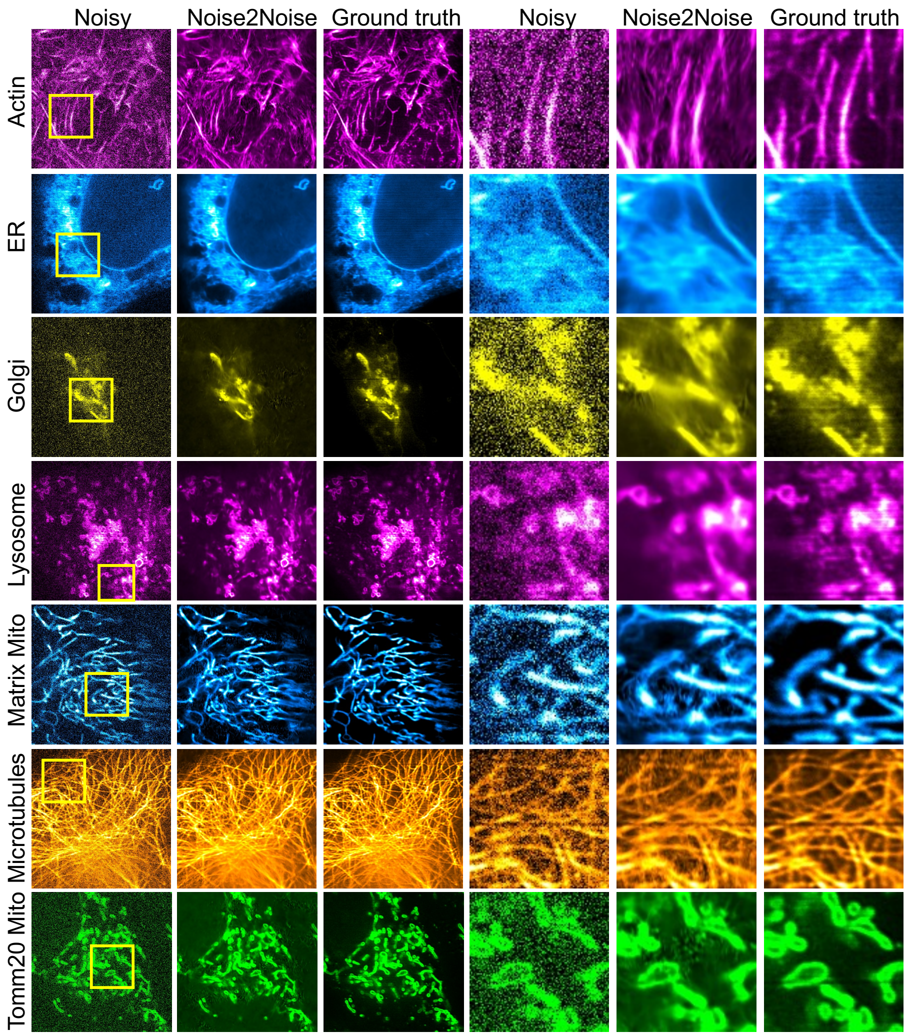


**Figure 9:** Qualitative results of the Noise2Noise plugin on a test sample from the 3D RCAN dataset [25]. From Left to right: noisy images (added Gaussian noise to the ground truth images), denoised images using Noise2Noise plugin, and target images (of deconvolution images in denoising folder from 3D RCAN dataset) as a single channel image, respectively. Selected ROI (in the yellow color region) for each image is shown on the right side in the same order, respectively. Scale bar, 5 $\mu$m.

modality lattice light-sheet microscopy (LLSM) system [26].

Quantitative results (PSNR and SSIM) of the fixed pattern noisy and Noise2Noise plugin denoised images for the HeLa microtubules and brine shrimp samples are provided in Table 3.

In addition, Figure 13(a) show the noisy dark-field microscopy image of

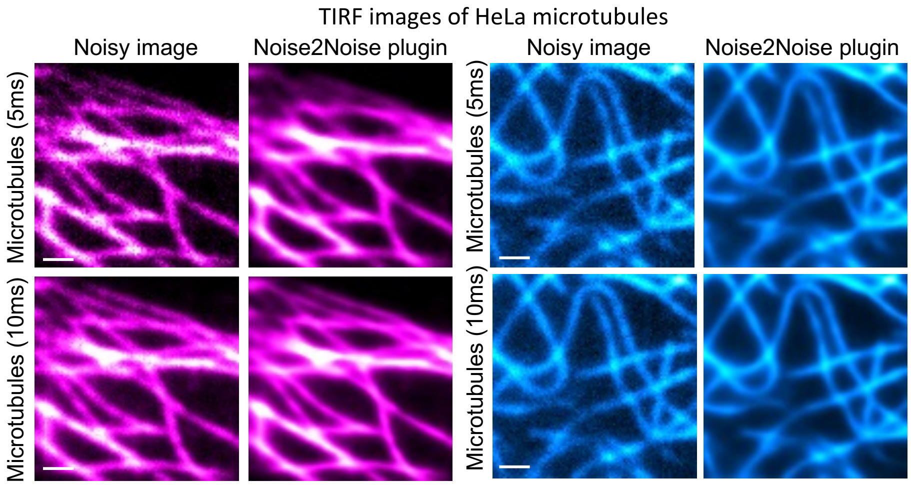


**Figure 10:** Qualitative results of the Noise2Noise plugin on a test sample from the ACsN dataset [26] where noisy images contain the fixed-pattern noise. From Left to right: noisy images and denoised images using Noise2Noise plugin as a single-channel image. Scale bar, 1 $\mu$m.

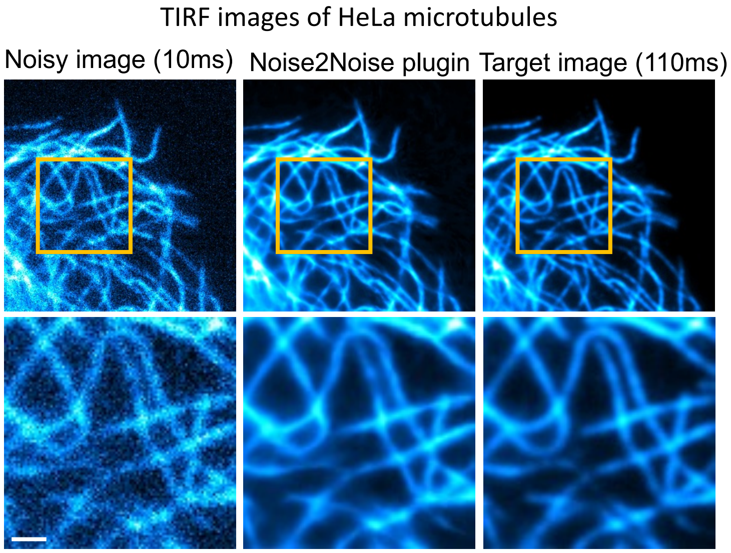


**Figure 11:** Qualitative results of the Noise2Noise plugin on a test sample from the ACsN dataset [26] where noisy images contain the fixed-pattern noise. From Left to right: noisy images, denoised images using Noise2Noise plugin, and target image a single channel image, respectively. Scale bar, 1 $\mu$m.

nanopores fabricated on a 300 nm thick gold-silicon dioxide ($SiO_2$) (200 nm

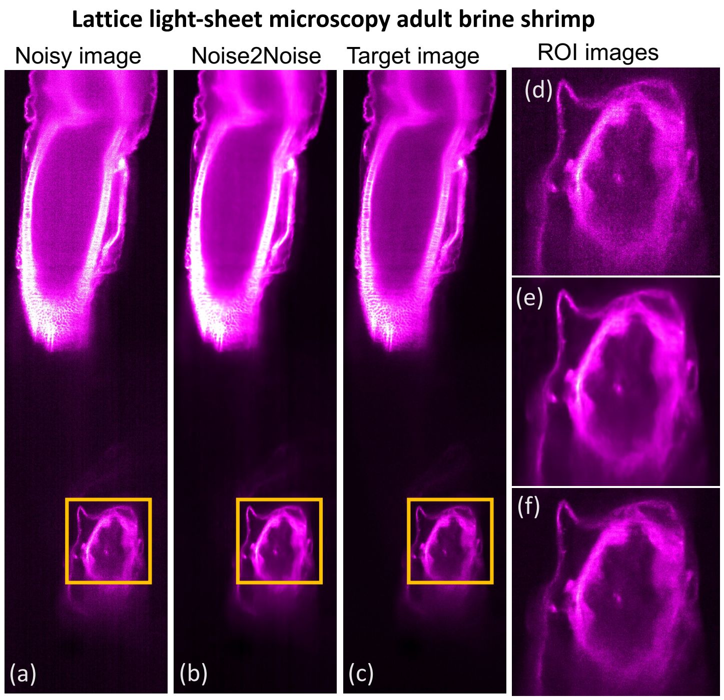


**Figure 12:** Qualitative results of the Noise2Noise plugin on a test sample from the ACsN dataset [26] where adult brine shrimp noisy images contain the fixed-pattern noise. From Left to right: noisy images (a, d), denoised images using Noise2Noise plugin (b, e), and target image (c, f) as a single channel image, respectively. Selected ROI (in the yellow color region) for each image is shown in the right most column (d, e, f respectively). Scale bar, 1 $\mu$m.

| **Sample** | **Raw/ Denoised** | **PSNR (dB)** | **SSIM** |
|---|---|---|---|
| Microtubules | Raw | 22.65 | 0.57 |
| Microtubules | Noise2Noise plugin | 29.13 | 0.83 |
| Brine shrimp | Raw | 25.44 | 0.56 |
| Brine shrimp | Noise2Noise plugin | 30.31 | 0.87 |

**Table 3:** Quantitative results PSNR and SSIM of our Noise2Noise plugin image denoising methods, when applied to the HeLa cell microtubules and brine shrimp samples with fixed pattern noise to the fluorescence microscopy images in the ACsN dataset [26].

gold, and 100 nm $SiO_2$) layer. Here, the pixel size is 150nm and the image dimension is 512 $\times$ 512 pixels. The denoised image by the Noise2Noise plugin is shown in Figure 13(b). The PSNR values of the noisy dark-field image and the denoised image are 21.17 dB and 30.16 dB, respectively. The target image

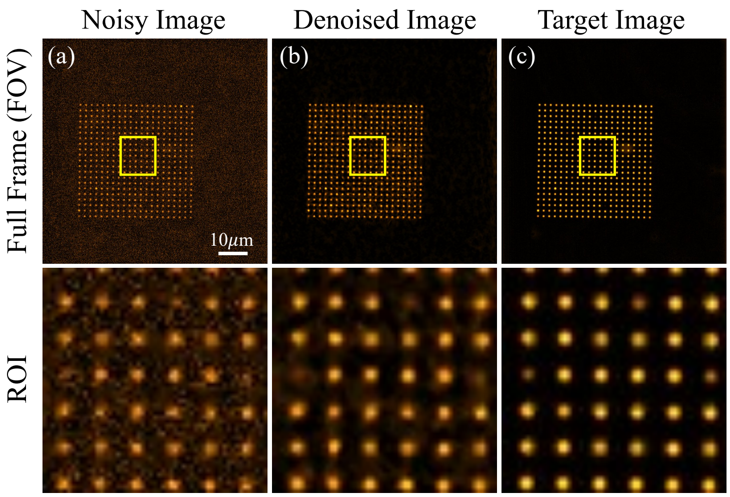


**Figure 13:** Noise2Noise plugin image denoising on a nanopores image captured using a dark-field microscope. (a) noisy image; (b) image denoised using Noise2Noise plugin and (c) target images, respectively. The full FOV has a size of 512 × 512 pixels. Selected ROI (in the yellow color region) for each image is shown in the bottom row.

is shown in Figure 13(c). Here the target image is the average of 500 noisy images within the same FOV. The denoising operation increases the PSNR by 8.99 dB. A small region of interest (ROI) is included in the bottom row for better visualization. Here, the denoised images have better quality and closely match with target images.

More fluorescence microscopy images and Noise2Noise plugin denoising results are provided in the GitHub repository[5]. From the above-mentioned test images of various noise levels, out-of-distribution structures, our demonstrated method (Noise2Noise plugin supervised ML model) is generalized since it is trained with the FMD dataset, which contains the mixture of Poisson-Gaussian noise on a large dataset includes fixed cells, in vivo samples. The illustrated results show the PSNR improvement of noisy images over a broad range of applications such as other fluorescence microscopy images.

### NOTE S5: ImageJ Plugin

Our Noise2Noise plugin ML model image denoising method is better than other existing methods and can provide better results for any fluorescence microscope noisy image. Also this new approach is based on deep convolutional neural network (with the Noise2Noise architecture) using noisy fluorescence microscopy images as input and requires significantly less computation time compared to

[5]`https://github.com/ND-HowardGroup/Instant-Image-Denoising/tree/master/Plugins/Model_validation`

alternative techniques. We observed there is a gap between the advanced image processing methods (using machine learning/convolutional neural networks) and easy usage of the image analysis tool (like ImageJ) by biologists, people in Optics/Biomedical image processing areas. Hence, we packaged our method as an ImageJ plugin that can denoise almost any microscopy image. We have developed the ImageJ plugins based on the ImageJ-tensorflow library [27]. Here the library provides the minimal set of functions in ImageJ to load the pre-trained ML model weights as a graph and perform a set of convolutional operations on the input image (loaded by ImageJ) which results in the output of the trained ML model.

We increased the impact of this novel work by reaching to more audience with easy access techniques. ImageJ plugins installation step by step procedure is provided [6]. The ImageJ denoising plugins JAR files and its corresponding source code in Java are available [7]. Figures shown in the main manuscript are generated with this ImageJ plugin and corresponding images are available [8]. For example, Figure 14 shows the noisy BPAE cell sample that is acquired with the commercial widefield microscope and its corresponding denoised images using the Noise2Noise plugin. In the same figure, the right most image indicates the ground-truth images (which is generated by averaging 50 noisy images with in the same FOV). Clearly the ML-based image denoising methods integrated with ImageJ shows the novelty of this work. Also we have updated the GITHUB repository with test data images in the read-me section which includes the qualitative and quantitative results.

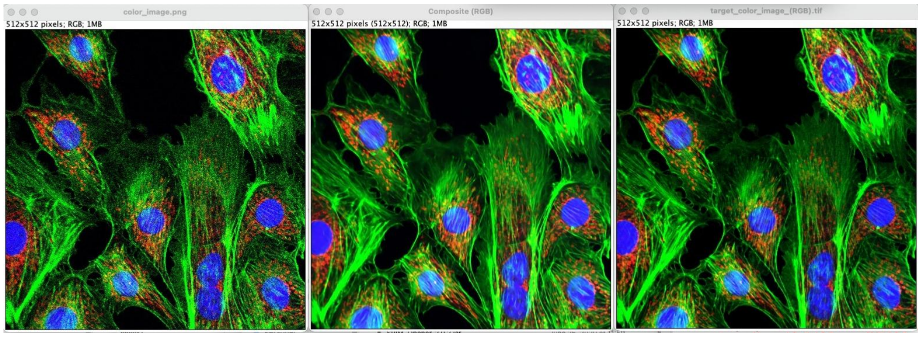


**Figure 14:** Noise2Noise plugin image denoising on a BPAE cell sample captured with a commercial confocal microscopy with pixel dwell time of 2 $\mu$s and the pixel width of 300 nm (from the FMD dataset). From Left to right: Noisy image, Denoised image using the Noise2Noise plugin and Target image respectively (the full-frame of size 512 $\times$ 512 pixels).

[6] https://github.com/ND-HowardGroup/Instant-Image-Denoising/blob/master/Plugins/Instructions_to_Install_Image_denoising_plugins.docx

[7] https://github.com/ND-HowardGroup/Instant-Image-Denoising/tree/master/Plugins/Image_Denoising_Plugins_Journal

[8] https://github.com/ND-HowardGroup/Instant-Image-Denoising/tree/master/Plugins/Test_images

#### Data and code availability

The FMD dataset mentioned in this paper is publicly available in the CurateND: `https://curate.nd.edu/show/f4752f78z6t`. The code for training the Noise2Noise plugin and DnCNN plugin architectures and underlying the results presented in the paper are publicly available in the GitHub repository: `https://github.com/ND-HowardGroup/Instant-Image-Denoising/`. Also, this repository includes the estimation of noise parameters using MATLAB codes, Noise2Noise plugin validation on W2S dataset, Noise2Noise plugin validation on out-of-distribution samples, and plugin source code in java with ImageJ plugins. Additional details of our Noise2Noise plugin denoising validation on the large datasets are available in CurateND: `https://curate.nd.edu/show/5h73pv66h5f`. Out-of-distribution structure dataset (3D RCAN dataset) used to generate underlying denoising results presented in this paper are available in Ref. [25].

#### Disclosures

The authors declare no conflicts of interest.

#### Funding Information

This material is based upon work supported by the National Science Foundation (NSF) under Grant No. CBET-1554516 and by the Department of Energy Office of Science under Grant DE-SC0019312.

#### Acknowledgments

Yide Zhang's research was supported by the Berry Family Foundation Graduate Fellowship of Advanced Diagnostics&Therapeutics (AD&T), University of Notre Dame. The authors acknowledge the Notre Dame Integrated Imaging Facility (NDIIF) for the use of the Nikon A1R-MP confocal microscope and Nikon Eclipse 90i widefield microscope in NDIIF's Optical Microscopy Core. The authors further acknowledge the Notre Dame Center for Research Computing (CRC) for providing the Nvidia GeForce GTX 1080-Ti GPU resources for training the Fluorescence Microscopy Denoising (FMD) dataset in Keras with TensorFlow backend.

[2] James Pawley. *Handbook of biological confocal microscopy*, volume 236. Springer Science & Business Media, 2006.

[3] Winfried Denk, James H Strickler, and Watt W Webb. Two-photon laser scanning fluorescence microscopy. *Science*, 248(4951):73–76, 1990.

[4] Peter J Verveer, Mark J Gemkow, and Thomas M Jovin. A comparison of image restoration approaches applied to three-dimensional confocal and wide-field fluorescence microscopy. *Journal of Microscopy*, 193(1):50–61, 1999.

[5] Varun Mannam, Yide Zhang, Yinhao Zhu, and Scott Howard. Fluorescence microscopy denoising (fmd) dataset, 2019.

[6] Yide Zhang, Yinhao Zhu, Evan Nichols, Qingfei Wang, Siyuan Zhang, Cody Smith, and Scott Howard. A poisson-gaussian denoising dataset with real fluorescence microscopy images. In *Proceedings of the IEEE Conference on Computer Vision and Pattern Recognition*, pages 11710–11718, 2019.

[7] Jaakko Lehtinen, Jacob Munkberg, Jon Hasselgren, Samuli Laine, Tero Karras, Miika Aittala, and Timo Aila. Noise2Noise: Learning image restoration without clean data. *arXiv preprint arXiv:1803.04189*, 2018.

[8] Francois Chollet et al. Keras, 2015.

[9] Diederik P Kingma and Jimmy Ba. Adam: A method for stochastic optimization. *arXiv preprint arXiv:1412.6980*, 2014.

[10] Leslie N Smith and Nicholay Topin. Super-convergence: Very fast training of neural networks using large learning rates. In *Artificial Intelligence and Machine Learning for Multi-Domain Operations Applications*, volume 11006, page 1100612. International Society for Optics and Photonics, 2019.

[19] Varun Mannam, Yide Zhang, Yinhao Zhu, and Scott Howard. Instant image denoising plugin for ImageJ using convolutional neural networks. In *Microscopy Histopathology and Analytics*, pages MW2A–3. Optical Society of America, 2020.
